# A hierarchy of cell death pathways confers layered resistance to shigellosis in mice

**DOI:** 10.1101/2022.09.21.508939

**Authors:** Justin L Roncaioli, Janet Peace Babirye, Roberto A Chavez, Fitty L Liu, Elizabeth A Turcotte, Angus Y Lee, Cammie F Lesser, Russell E Vance

## Abstract

Bacteria of the genus *Shigella* cause shigellosis, a severe gastrointestinal disease driven by bacterial colonization of colonic intestinal epithelial cells. Vertebrates have evolved programmed cell death pathways that sense invasive enteric pathogens and eliminate their intracellular niche. Previously we reported that genetic removal of one such pathway, the NAIP–NLRC4 inflammasome, is sufficient to convert mice from resistant to susceptible to oral *Shigella flexneri* challenge (Mitchell, Roncaioli et al., 2020). Here, we investigate the protective role of additional cell death pathways during oral mouse *Shigella* infection. We find that the Caspase-11 inflammasome, which senses *Shigella* LPS, restricts *Shigella* colonization of the intestinal epithelium in the absence of NAIP–NLRC4. However, this protection is limited when *Shigella* expresses OspC3, an effector that antagonizes Caspase-11 activity. TNFα, a cytokine that activates Caspase-8-dependent apoptosis, also provides protection from *Shigella* colonization of the intestinal epithelium, but only in the absence of both NAIP– NLRC4 and Caspase-11. The combined genetic removal of Caspases-1,-11, and -8 renders mice hyper-susceptible to oral *Shigella* infection. Our findings uncover a layered hierarchy of cell death pathways that limit the ability of an invasive gastrointestinal pathogen to cause disease.

## Introduction

*Shigella* is a genus of enteric bacterial pathogens that causes ∼270 million yearly cases of shigellosis, with ∼200,000 of these resulting in death (Khalil et al., 2018). Shigellosis manifests as an acute inflammatory colitis resulting in abdominal cramping, fever, and in severe cases, bloody diarrhea (dysentery) (Kotloff et al., 2018). Bacterial invasion of the colonic intestinal epithelium and subsequent dissemination between adjacent intestinal epithelial cells (IECs) is believed to drive inflammation and disease. *Shigella* pathogenesis is mediated by a virulence plasmid which encodes a type three secretion system (T3SS) and more than 30 virulence factors or effectors (Schnupf and Sansonetti, 2019; Schroeder and Hilbi, 2008). The T3SS injects effectors into the host cell that facilitate bacterial uptake, escape into the cytosol, and actin–based motility, which is essential for bacterial spread to neighboring epithelial cells (Bernardini et al., 1989; Goldberg and Theriot, 1995; Mattock and Blocker, 2017). The T3SS also secretes several effectors that disarm the host innate immune response to make the cell a more hospitable niche for replicating *Shigella* (Ashida et al., 2015).

The innate immune system can counteract intracellular bacterial pathogens by inducing programmed cell death (Williams, 1994). Programmed cell death eliminates the intracellular pathogen niche, maintains epithelial barrier integrity, promotes clearance of damaged cells, and enhances presentation of foreign antigens to cells of the adaptive immune response (Deets et al., 2021; Doran et al., 2020; Jorgensen et al., 2017; Koch and Nusrat, 2012; Yatim et al., 2017). Three main modes of programmed cell death are common to mammalian cells: pyroptosis, apoptosis, and necroptosis. Each is controlled by distinct sensors and conserved downstream executors which together provide a formidable barrier that pathogens must subvert for successful intracellular replication. Of particular relevance to *Shigella* and other gastrointestinal pathogens, cell death of IECs is accompanied by a unique cellular expulsion process that rapidly and selectively ejects dying or infected cells from the epithelial layer, thereby potently limiting pathogen invasion into deeper tissue (Fattinger et al., 2021; Knodler et al., 2014; Rauch et al., 2017; Sellin et al., 2014).

*Shigella* is an example of a pathogen in intense conflict with host cell death pathways (Ashida et al., 2021). *Shigella* encodes multiple effectors to prevent cell death in human cells, including OspC3 to block Caspase-4 inflammasome activation (Kobayashi et al., 2013; Li et al., 2021; Mou et al., 2018; Oh et al., 2021), IpaH7.8 to inhibit Gasdermin D-dependent pyroptosis (Luchetti et al., 2021), OspC1 to suppress Caspase-8-dependent apoptosis (Ashida et al., 2020), and OspD3 to block necroptosis (Ashida et al., 2020). The antagonism of these pathways (and perhaps others that are yet undiscovered) and the resulting maintenance of the epithelial niche appears sufficient to render humans susceptible to *Shigella* infection. Mice, however, are resistant to oral *Shigella* challenge because *Shigella* is unable to counteract epithelial NAIP–NLRC4-dependent cell death and expulsion (Chang et al., 2013; Mitchell, Roncaioli et al., 2020). Removal of the NAIP–NLRC4 inflammasome renders mice susceptible to shigellosis, providing a tractable genetic model to dissect *Shigella* pathogenesis after oral infection *in vivo* (Mitchell, Roncaioli et al., 2020).

Here, we use the NAIP–NLRC4-deficient mouse model of shigellosis to investigate the role of programmed cell death in defense against *Shigella in vivo*. We find that Caspase-11 (CASP11), a cytosolic sensor of LPS and the mouse ortholog of Caspase-4 (Shi et al., 2014), provides modest protection from *Shigella* infection in the absence of NAIP–NLRC4. Like in humans, this pathway is antagonized by the *Shigella* effector OspC3, and genetic removal of *ospC3* from *Shigella* results in a significant reduction in bacterial colonization of IECs and virulence which depends on CASP11. We also find that TNFα, a cytokine that can induce TNF receptor 1 (TNFRI)-dependent extrinsic apoptosis (Piguet et al., 1998), defends mouse IECs from bacterial colonization and limits subsequent disease. TNFα-dependent protection, however, was only observed when mice lack both NAIP–NLRC4 and CASP11, revealing a hierarchal program of cell death pathways that counteract *Shigella in vivo. Casp1/11*^*–/–*^*Ripk3*^*–/–*^ and *Casp8*^*–/–*^*Ripk3*^*–/–*^ mice, which lack some but not all key components of pyroptosis, apoptosis, and necroptosis are largely protected from disease, revealing redundancies between these pathways. *Casp1/11/8*^*–/–*^*Ripk3*^*–/–*^ mice, however, are hyper-susceptible to shigellosis, indicating that programmed cell death is a predominant host defense mechanism against *Shigella* infection. Furthermore, neither Interleukin-1 receptor (IL-1R)-mediated signaling nor myeloid-restricted NAIP–NLRC4 have an apparent effect on *Shigella* pathogenesis, suggesting that it is cell death of IECs that primarily protects mice from shigellosis. Our findings underscore the importance of cell death in defense against intracellular bacterial pathogens and provide an example of how layered and hierarchical immune pathways can provide robust defense against pathogens that have evolved a broad arsenal of virulence factors.

## Results

### CASP11 contributes to resistance of B6 versus 129 *Nlrc4*^*–/–*^ mice to shigellosis

We previously generated NLRC4 deficient mice on the 129S1/SvImJ (129) background (129.*Nlrc4*^*–/–*^) and observed that these mice appeared more susceptible to oral *Shigella flexneri* challenge than C57BL/6J (B6) NLRC4 deficient mice (B6.*Nlrc4*^*–/–*^) (Mitchell, Roncaioli et al., 2020). We reasoned that the apparent difference between the strains might be due to genetic and/or microbiota differences. To address these possibilities, we infected co-housed B6.*Nlrc4*^*–/–*^ and 129.*Nlrc4*^*–/–*^ mice and directly compared disease severity between the two strains (Figure 1). The B6.*Nlrc4*^*–/–*^ mice exhibited only modest weight-loss (5-10% of starting weight) through two days and began to recover by day three (Figure 1A). The 129.*Nlrc4*^*–/–*^ mice, however, continued to lose weight through day three (10-15% of starting weight) (Figure 1A). Upon sacrifice at day three, we harvested the intestinal epithelial cell fraction from the cecum and colon of each mouse, washed this fraction in gentamicin to eliminate any extracellular *Shigella*, and lysed these cells to enumerate intracellular bacterial colonization of IECs. IECs from 129.*Nlrc4*^*–/–*^ mice harbored >10-fold higher intracellular *Shigella* burdens than those from B6.*Nlrc4*^*–/–*^ mice (Figure 1B). We also found that 129.*Nlrc4*^*–/–*^ mice had higher levels of inflammatory cytokines CXCL1 and IL-1β in their intestinal tissue, as measured by ELISA (Figure 1C, D) and exhibited significantly more gross cecum shrinkage than B6.*Nlrc4*^*–/–*^ mice (Figure 1E). The 129.*Nlrc4*^*–/–*^ mice also exhibited more pronounced diarrhea (as measured by the wet weight to dry weight ratio of mouse feces) relative to the B6.*Nlrc4*^*–/–*^ mice at two days post-infection (Figure 1F). We scored mouse feces for the presence of occult blood (score = 1) or macroscopic blood (score = 2) at day two and three, the sum of which represents a blood score from zero to four (Figure 1G). All 129.*Nlrc4*^*–/–*^ mice had occult blood in their feces on at least one of these days, with many having occult or macroscopic blood on both days. In contrast, B6.*Nlrc4*^*–/–*^ mice did not exhibit fecal blood.

**Figure 1.**
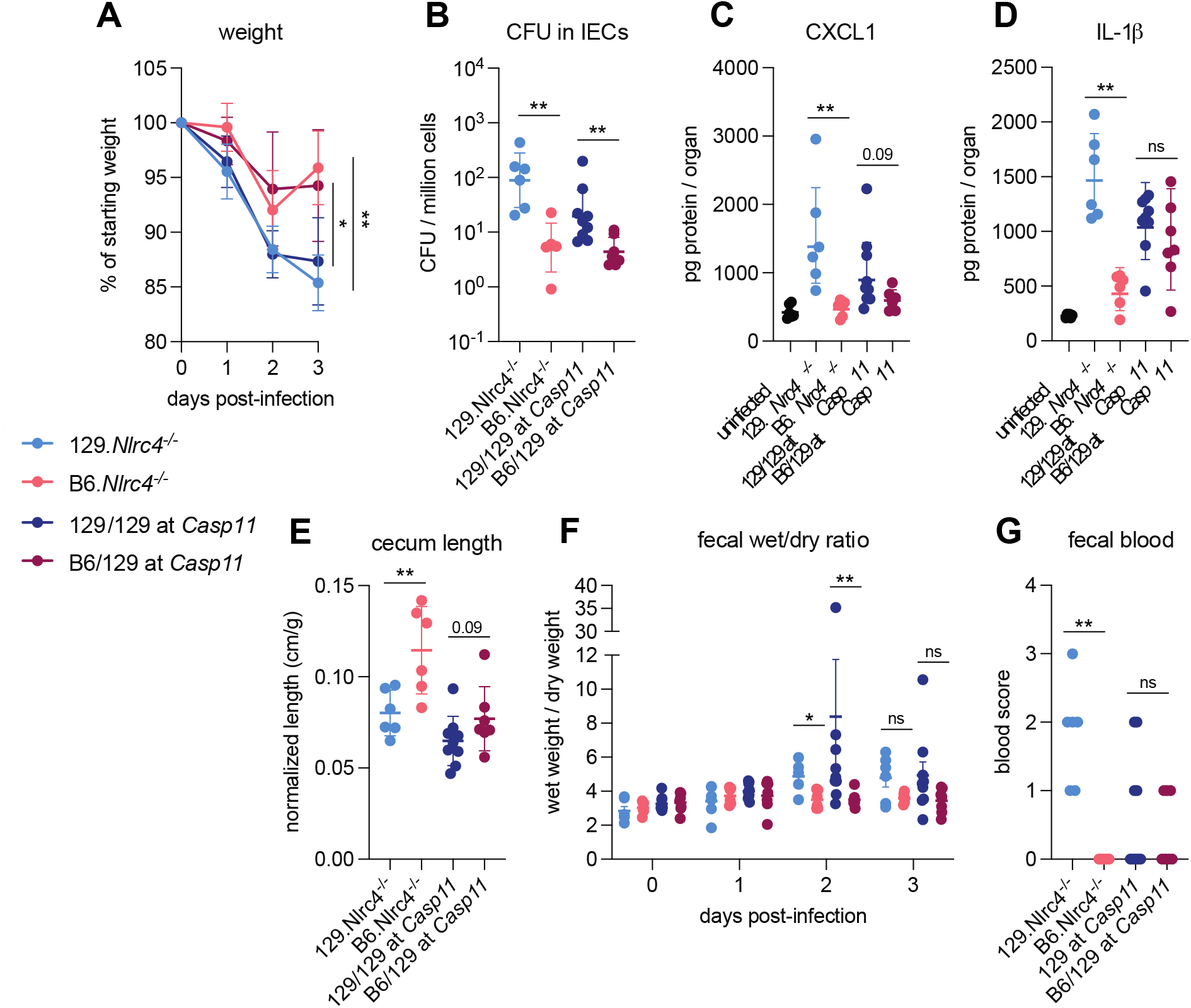
CASP11 contributes to resistance of B6 versus 129 *Nlrc4*^*–/–*^ mice to shigellosis. (**A-G**) B6.*Nlrc4*^*–/–*^ (pink), 129.*Nlrc4*^*–/–*^ (light blue), backcrossed littermates that are homozygous 129/129 at *Casp11* (dark blue), and backcrossed littermates that are heterozygous B6/129 at *Casp11* (maroon) were co-housed for 3 weeks, treated orally with 25 mg streptomycin sulfate in water, and orally challenged the next day with 10^7^ CFU of WT *Shigella flexneri*. Mice were sacrificed at three days post-infection. (**A**) Mouse weights from 0 through 3 days post-infection. Each symbol represents the mean for all mice of the indicated genotype. (**B**) *Shigella* colony forming units (CFU) per million cells from the combined intestinal epithelial cell (IEC) enriched fraction of gentamicin-treated cecum and colon tissue. (**C, D**) CXCL1 and IL-1β levels measured by ELISA from homogenized cecum and colon tissue of infected mice. (**E**) Quantification of cecum lengths normalized to mouse weight prior to infection; cecum length (cm) / mouse weight (g). (**F**) The ratio of fecal pellet weight when wet (fresh) divided by the fecal pellet weight after overnight drying. A larger wet/dry ratio indicates increased diarrhea. Pellets were collected daily from 0-3 days post-infection. (**G**) Additive blood scores from feces collected at two and three days post-infection. 1 = occult blood, 2 = macroscopic blood for a given day, maximum score is 4. (**B-G**) Each symbol represents one mouse. Mean ± SD is shown in (**A, C-E**). Geometric mean ± SD is shown in (**B**). Mean ± SEM is shown in **(F**). Mann-Whitney test, *p < 0.05, **p < 0.01, ***p < 0.001, ****p < 0.0001, ns = not significant (p > 0.05).

The persistence of this difference in disease severity between co-housed 129 and B6 *Nlrc4*^−/–^ mice suggested that genetic rather than microbiota differences might explain the differential susceptibility of the strains. The mouse non-canonical inflammasome Caspase-11 (CASP11) and its human orthologs Caspases-4 and -5 sense cytosolic *Shigella* LPS to initiate pyroptosis (Hagar et al., 2013; Kayagaki et al., 2011; Kobayashi et al., 2013; Shi et al., 2014). Notably, 129 mice are naturally deficient for Caspase-11 (Kayagaki et al., 2011). To determine if Caspase-11 contributes to the difference in susceptibility between these strains, we crossed B6.*Nlrc4*^*–/–*^ and 129.*Nlrc4*^*–/–*^ mice to generate B6/129.*Nlrc4*^*–/–*^ F_1_ hybrids (Figure 1 — figure supplement 1A). We infected these F_1_ hybrids and found that they were relatively resistant to *Shigella* challenge and resembled the parental B6.*Nlrc4*^*–/–*^ mice (Figure 1 — figure supplement 1B-F). These results are consistent with the possibility that a dominant gene on the C57BL/6J background provides protection from *Shigella*. Next, we backcrossed these hybrids to the 129.*Nlrc4*^*–/–*^ parental strain to generate littermate *Nlrc4*^*–/–*^ mice that were mixed homozygous 129/129 or heterozygous B6/129 at all loci (Figure 1 — figure supplement 1A). We co-housed these *Nlrc4*^*–/–*^ backcrossed mice with their parental 129.*Nlrc4*^*–/–*^ and B6.*Nlrc4*^*–/–*^ strains for >3 weeks, infected them with *Shigella*, and genotyped each at the *Casp11* locus to determine whether a functional B6 *Casp11* allele would correlate with reduced disease severity.

Indeed, backcrossed *Nlrc4*^*–/–*^ mice that were heterozygous B6/129 at *Casp11* were more resistant to shigellosis than backcrossed *Nlrc4*^*–/–*^ mice that were 129/129 at *Casp11* (Figure 1). Mice that were heterozygous B6/129 at *Casp11* showed a similar weight-loss pattern to the parental B6.*Nlrc4*^*–/–*^ mice and began to recover by day three while the weight-loss in mice that were homozygous 129/129 at *Casp11* phenocopied that of the parental 129.*Nlrc4*^*–/–*^ mice (Figure 1A). Consistent with these results, mice that were homozygous 129/129 at *Casp11* exhibited enhanced bacterial colonization of the intestinal epithelium (Figure 1B), modest increases in inflammatory cytokines (Figure 1C, D) and cecum shrinkage (Figure 1E), and more pronounced diarrhea (Figure 1F). Despite these differences, there was no strong correlation between fecal blood score and *Casp11* genotype (Figure 1G), suggesting that while *Casp11* contributes to resistance, there are additional genetic modifiers present on the 129 or B6 background that affect susceptibility to shigellosis. As these additional modifiers appear to be relatively weak compared to *Casp11*, we did not attempt to map them genetically.

However, we did specifically test for a contribution of *Hiccs*, a genetic locus in 129 mice that associates with increased susceptibility to *Helicobacter hepaticus*-dependent colitis (Boulard et al., 2012). In contrast to *Casp11*, we found that *Hiccs* did not significantly associate with increased susceptibility to shigellosis in 129 mice (Figure 1 — figure supplement 2).

### CASP11 prevents IEC colonization and disease in B6.*Nlrc4*^*–/–*^ mice

To define the role of mouse Caspase-11 in a uniform genetic background, we generated *Casp11*^***–/–***^ mice on the B6.*Nlrc4*^*–/–*^ background using CRISPR-Cas9 editing (Figure 2 — figure supplement 1). We previously found that *Casp1/11*^*–/–*^ mice are largely resistant to oral WT *Shigella flexneri* infection, likely because NLRC4-dependent Caspase-8 activation is sufficient to prevent bacterial colonization of IECs (Figure 2 — figure supplement 2) (Mitchell, Roncaioli et al., 2020; Rauch et al., 2017). Thus, Caspase-11 is dispensable for protection from WT *Shigella* challenge when mice express functional NLRC4, but Caspase-11 could still be critical as a backup pathway in the absence of NLRC4. We therefore challenged B6.*Nlrc4*^*–/–*^ and B6.*Nlrc4*^*–/–*^ *Casp11*^*–/–*^ littermates with WT *Shigella* and assessed pathogenicity for two days following infection.

We observed a modest increase in susceptibility to *Shigella* infection in B6.*Nlrc4*^*–/–*^*Casp11*^*–/–*^ mice relative to B6.*Nlrc4*^*–/–*^ mice (Figure 2). While B6.*Nlrc4*^*–/–*^*Casp11*^*–/–*^ mice did not experience more weight loss (Figure 2A), cecum shrinkage (Figure 2B), or diarrhea (Figure 2C) than B6.*Nlrc4*^*–/–*^ mice, there was a 5-fold increase in *Shigella* burdens in IECs from B6.*Nlrc4*^*–/–*^*Casp11*^*–/–*^ mice (Figure 2D), indicating that Caspase-11 protects the mouse epithelium from bacterial colonization in the absence of NLRC4. Intestinal tissue from B6.*Nlrc4*^*–/–*^*Casp11*^*–/–*^ mice also expressed higher levels of inflammatory cytokines CXCL1 and IL-1β than tissue from B6.*Nlrc4*^*–/–*^ mice (Figure 2E, F). Consistent with these findings, B6.*Nlrc4*^*–/–*^ mice did not exhibit blood in their feces but two of the nine B6.*Nlrc4*^*–/–*^*Casp11*^*–/–*^ did present with occult blood (Figure 2G). These results suggest that Caspase-11 has a relatively modest contribution to defense against wild-type *Shigella*. Indeed, a minor role for Caspase-11 is expected given that *Shigella* is known to encode an effector called OspC3 that inhibits Caspase-11 (see below). Nevertheless, taken together, our results in mixed 129/B6.*Nlrc4*^*–/–*^ and B6.*Nlrc4*^*–/–*^*Casp11*^*–/–*^ mice indicate that Caspase-11 contributes to defense against *Shigella in vivo* as a backup pathway in the absence of NLRC4 (Figure 2 — figure supplement 2).

**Figure 2.**
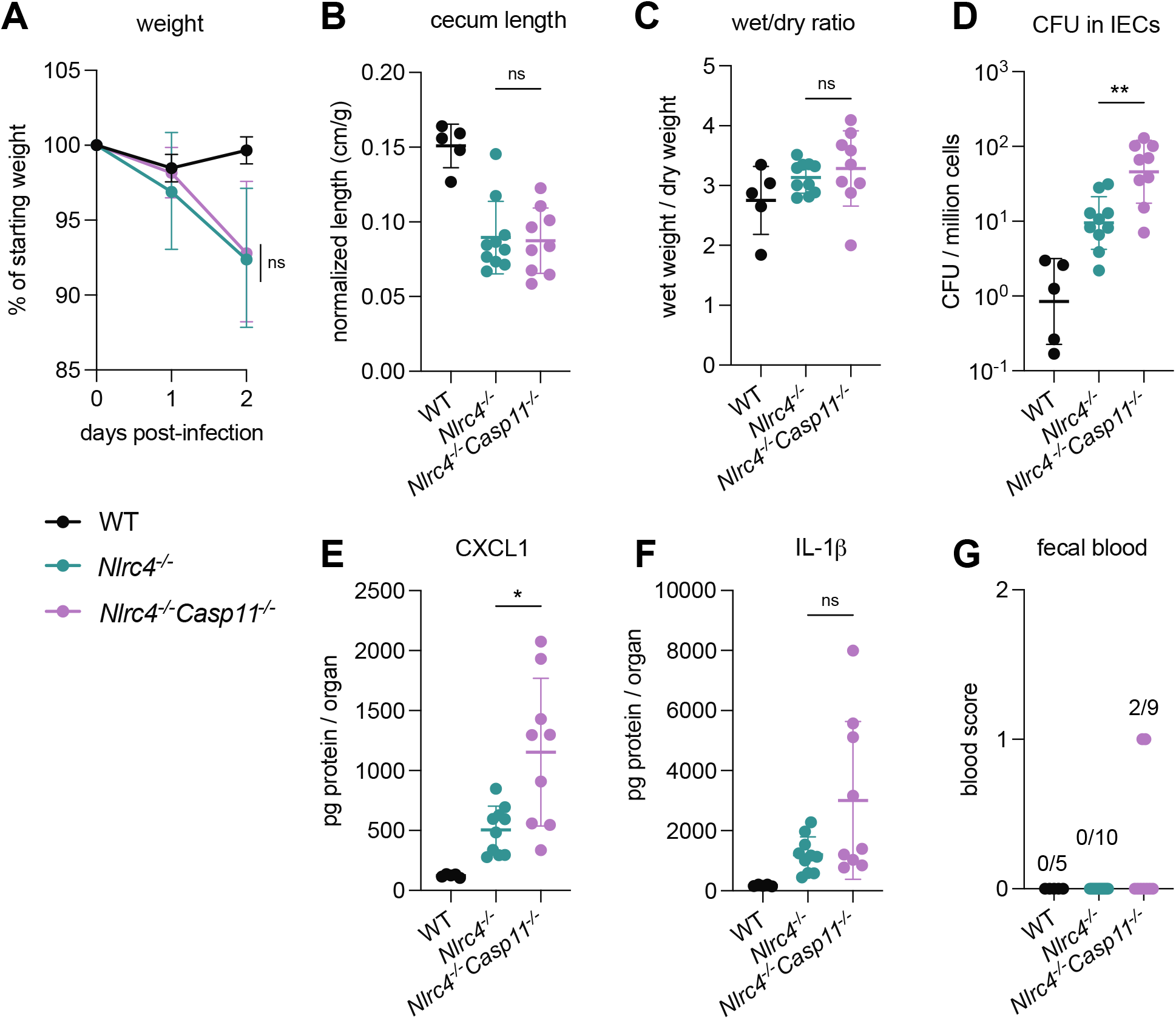
CASP11 prevents IEC colonization and disease in B6.*Nlrc4*^*–/–*^ mice. (**A**-**G**) B6.WT (black), B6.*Nlrc4*^*–/–*^ (teal), and B6.*Nlrc4*^*–/–*^*Casp11*^*–/–*^ (lavender) littermates were treated orally with 25 mg streptomycin sulfate in water and orally challenged the next day with 10^7^ CFU of WT *Shigella flexneri*. Mice were sacrificed at two days post-infection. (**A**) Mouse weights from 0 through 2 days post-infection. Each symbol represents the mean for all mice of the indicated genotype. (**B**) Quantification of cecum lengths normalized to mouse weight prior to infection; cecum length (cm) / mouse weight (g). (**C**) The ratio of fecal pellet weight when wet (fresh) divided by the fecal pellet weight after overnight drying. Pellets were collected at day two post-infection. A larger wet/dry ratio indicates increased diarrhea. (**D**) *Shigella* colony forming units (CFU) per million cells from the combined intestinal epithelial cell (IEC) enriched fraction of gentamicin-treated cecum and colon tissue. (**E, F**) CXCL1 and IL-1β levels measured by ELISA from homogenized cecum and colon tissue of infected mice. (**G**) Blood scores from feces collected at two days post-infection. 1 = occult blood, 2 = macroscopic blood. (**B-G**) Each symbol represents one mouse. Data collected from two independent experiments. Mean ± SD is shown in (**A-C, E, F**). Geometric mean ± SD is shown in (**D**). Mann-Whitney test, *p < 0.05, **p < 0.01, ***p < 0.001, ****p < 0.0001, ns = not significant (p > 0.05).

### *Shigella* effector OspC3 is critical for virulence in oral *Shigella* infection

*Shigella flexneri* protein OspC3 is a T3SS-secreted effector that inhibits both human Caspase-4 and mouse Caspase-11 to suppress pyroptosis (Kobayashi et al., 2013; Li et al., 2021; Mou et al., 2018; Oh et al., 2021). While OspC3 has been shown to be required for virulence during intraperitoneal mouse infection (Li et al., 2021; Oh et al., 2021) and for intestinal lumenal colonization in wild-type mice (Alphonse et al., 2022), the role of this effector has not been studied in an oral mouse model of infection where *Shigella* invades and replicates within the intestinal epithelium. Indeed, our results indicating a role for Caspase-11 in defense against wild-type *Shigella* (see above, Figures 1 and 2) suggested that the inhibition of Caspase-11 by OspC3 could be incomplete in epithelial cells. To test the role of OspC3 in shigellosis, we orally infected B6.*Nlrc4*^*–/–*^ mice with WT S*higella flexneri* or a mutant stain that lacks OspC3 (*ΔospC3*) (Figure 3). Consistent with our previous experiments, B6.*Nlrc4*^*–/–*^ mice challenged with WT S*higella* developed shigellosis characterized by weight-loss, increases in bacterial colonization of the intestinal epithelium, cecum shrinkage, diarrhea, and inflammatory cytokines (Figure 3A-G). However, B6.*Nlrc4*^*–/–*^ mice challenged with *ΔospC3 Shigella flexneri* were almost fully resistant to infection (Figure 3). *ΔospC3*-infected B6.*Nlrc4*^*–/–*^ mice still experienced some weight-loss (Figure 3B) and cecum shrinkage (Figure 3A, D), but exhibited a >10-fold decrease in IEC colonization (Figure 3C) and reduced levels of inflammatory cytokines relative to WT-infected B6.*Nlrc4*^*–/–*^ mice (Figure 3F, G).

**Figure 3.**
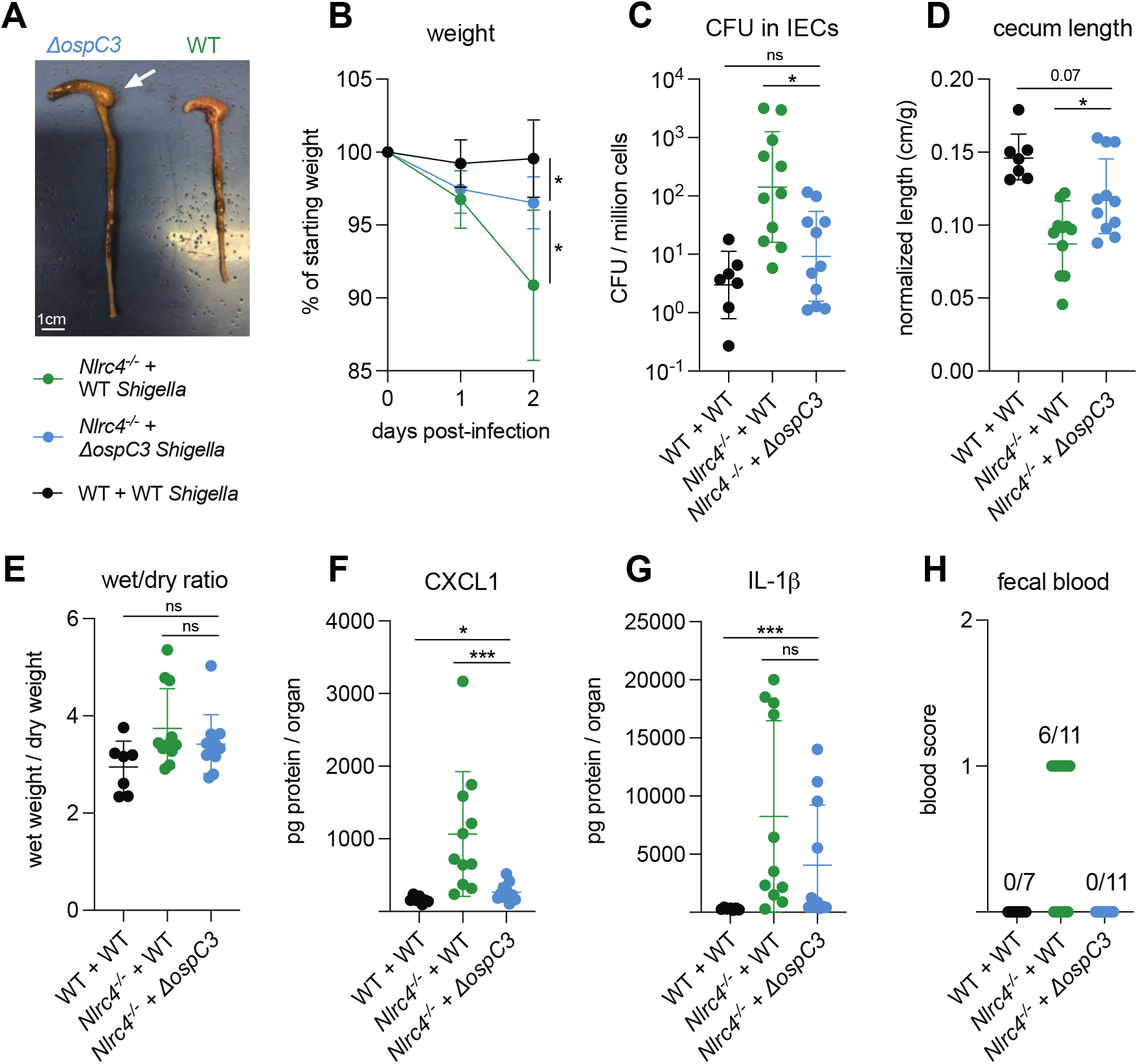
*Shigella* effector OspC3 is critical for virulence in oral *Shigella* infection. (**A-H**) Mice were treated orally with 25 mg streptomycin sulfate in water and infected one day later. B6.WT mice were orally challenged with 10^7^ CFU of WT *Shigella flexneri* (black) and B6.*Nlrc4*^***–/–***^ co-housed/littermate mice were challenged with WT (green) or *ΔospC3 Shigella flexneri* (blue). Mice were sacrificed at two days post-infection. (**A**) Representative images of the cecum and colon from B6.*Nlrc4*^***–/–***^ mice infected with WT or *ΔospC3 Shigella flexneri*. The white arrow indicates clear but reduced inflammation in mice infected with the *ΔospC3* strain. (**B**) Mouse weights from 0 through 2 days post-infection. Each symbol represents the mean for all mice of the indicated genotype. (**C**) *Shigella* colony forming units (CFU) per million cells from the combined intestinal epithelial cell (IEC) enriched fraction of gentamicin-treated cecum and colon tissue. (**D**) Quantification of cecum lengths normalized to mouse weight prior to infection; cecum length (cm) / mouse weight (g). (**E**) The ratio of fecal pellet weight when wet (fresh) divided by the fecal pellet weight after overnight drying. Pellets were collected at day two post-infection. (**F, G**) CXCL1 and IL-1β levels measured by ELISA from homogenized cecum and colon tissue of infected mice. (**H**) Blood scores from feces collected at two days post-infection. 1 = occult blood, 2 = macroscopic blood. (**C-H**) Each symbol represents one mouse. Data collected from two independent experiments. Mean ± SD is shown in (**B, D-G**). Geometric mean ± SD is shown in (**C**). Mann-Whitney test, *p < 0.05, **p < 0.01, ***p < 0.001, ****p < 0.0001, ns = not significant (p > 0.05).

Importantly, B6.*Nlrc4*^*–/–*^ mice infected with *ΔospC3 Shigella flexneri* did not display fecal blood while, in this experiment, six of the eleven B6.*Nlrc4*^*–/–*^ mice infected with WT *Shigella* did present with fecal blood (Figure 3H). These results indicate that *ΔospC3 Shigella* is significantly attenuated in our B6.*Nlrc4*^*–/–*^ mouse model of shigellosis.

OspC3 directly inactivates mouse Caspase-11 (Li et al., 2021) but has also been reported to modulate other signaling pathways, including interferon signaling (Alphonse et al., 2022). To test if the effect of OspC3 on virulence is dependent on inhibition of mouse Caspase-11, we infected both B6.*Nlrc4*^*–/–*^ and B6.*Nlrc4*^*–/–*^ *Casp11*^*–/–*^ mice with either WT or *ΔospC3 Shigella* strains. We again observed that the *ospC3* mutant was attenuated relative to WT *Shigella* in B6.*Nlrc4*^*–/–*^ mice (Figure 4). However, both WT and *ΔospC3 Shigella* caused severe disease in B6.*Nlrc4*^*–/–*^*Casp11*^*–/–*^ mice, with comparable weight-loss, bacterial colonization of the intestinal epithelium, cecum lengths, diarrhea, and fecal blood (Figure 4A-D, G). These results suggest that Caspase-11 is the primary physiological target of OspC3 *in vivo*. We did observe a significant increase in inflammatory cytokines in WT-infected B6.*Nlrc4*^*–/–*^*Casp11*^*–/–*^ mice relative to *ΔospC3*-infected B6.*Nlrc4*^*–/–*^ *Casp11*^*–/–*^ mice (Figure 4E, F), indicating that OspC3 might also affect immune pathways independent of Caspase-11. Again, we only observed a modest difference in disease severity between B6.*Nlrc4*^*–/–*^ and B6.*Nlrc4*^*–/–*^*Casp11*^*–/–*^ mice infected with WT *Shigella* (Figure 4), consistent with the ability of OspC3 to significantly reduce Caspase-11 activity. These results confirm prior reports that OspC3 inhibits Caspase-11 *in vivo* (Li et al., 2021; Oh et al., 2021) and further show that OspC3-dependent inhibition of Caspase-11 is required for *Shigella* virulence. Nonetheless, this inhibition is likely incomplete, as Caspase-11 still provides a degree of protection in B6.*Nlrc4*^*–/–*^ mice even when *Shigella* expresses OspC3 (Figures 2, 4).

**Figure 4.**
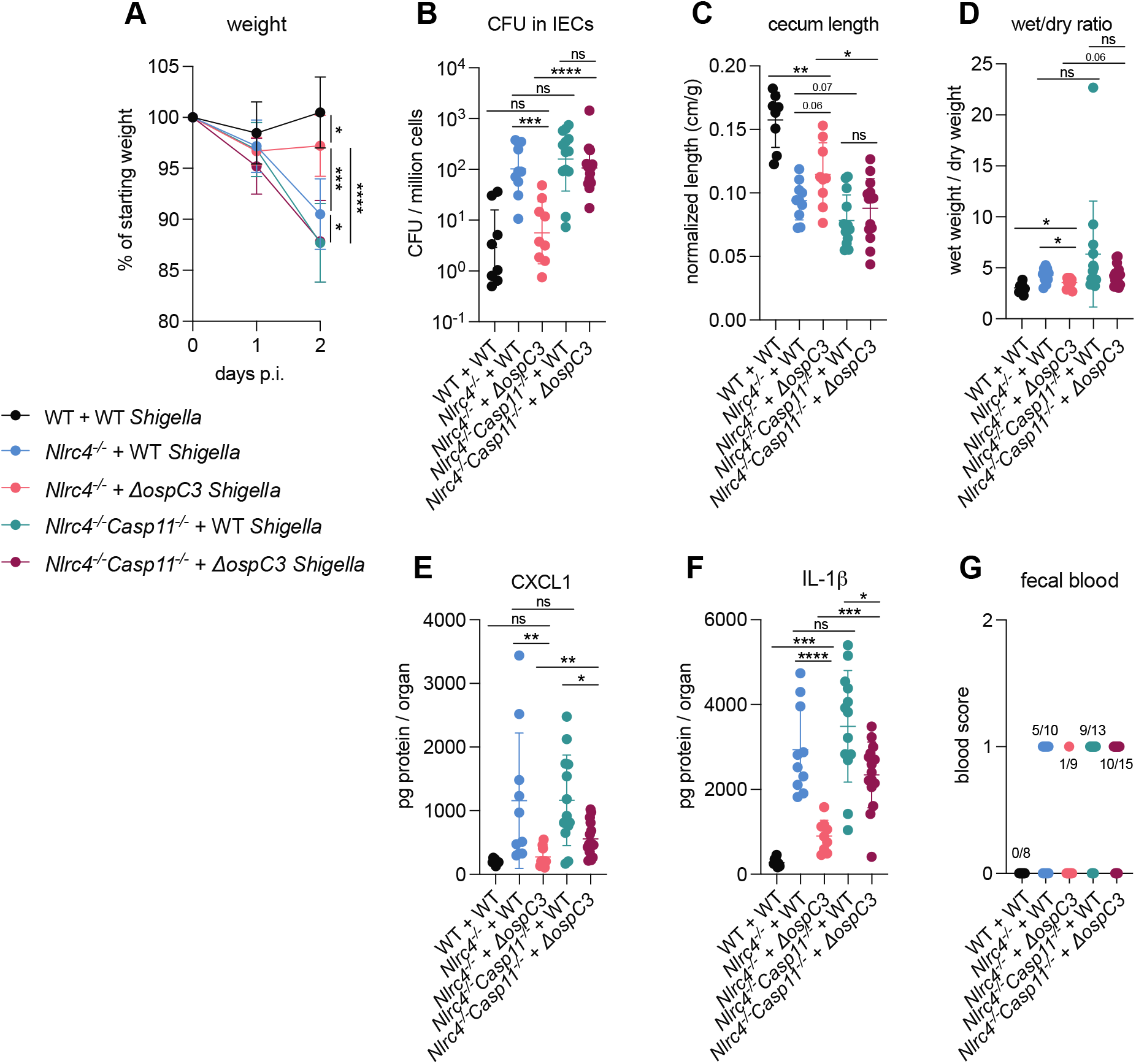
OspC3-driven virulence in B6.*Nlrc4*^*–/–*^ mice depends on Caspase-11. (**A-G**) Mice were treated orally with 25 mg streptomycin sulfate in water and then infected one day later. B6.WT mice were orally challenged with 10^7^ CFU of WT *Shigella flexneri* (black), B6.*Nlrc4*^***–/–***^ mice were challenged with WT (blue) or *ΔospC3 Shigella flexneri* (pink), and B6.*Nlrc4*^***–/–***^*Casp11*^***–/–***^ mice were challenged with WT (teal) or *ΔospC3 Shigella flexneri* (maroon). Mice were littermates or were co-housed for 3 weeks prior to infection and were sacrificed at two days post-infection. (**A)** Mouse weights from 0 through 2 days post-infection. Each symbol represents the mean for all mice of the indicated group. (**B**) *Shigella* colony forming units (CFU) per million cells from the combined intestinal epithelial cell (IEC) enriched fraction of gentamicin-treated cecum and colon tissue. (**C**) Quantification of cecum lengths normalized to mouse weight prior to infection; cecum length (cm) / mouse weight (g). (**D**) The ratio of fecal pellet weight when wet (fresh) divided by the fecal pellet weight after overnight drying. Pellets were collected at day two post-infection. (**E, F**) CXCL1 and IL-1β levels measured by ELISA from homogenized cecum and colon tissue of infected mice. (**G**) Blood scores from feces collected at two days post-infection. 1 = occult blood, 2 = macroscopic blood. (**B-G**) Each symbol represents one mouse. Data collected from two independent experiments. Mean ± SD is shown in (**A, C-F**). Geometric mean ± SD is shown in (**B**). Mann-Whitney test, *p < 0.05, **p < 0.01, ***p < 0.001, ****p < 0.0001, ns = not significant (p > 0.05).

### Neither myeloid NLRC4 nor IL-1 affect *Shigella* pathogenesis

The generally accepted model of *Shigella* pathogenesis proposes that *Shigella* cross the colonic epithelium via transcytosis through M-cells (Schnupf and Sansonetti, 2019; Schroeder and Hilbi, 2008). After transcytosis, *Shigella* is then believed to be phagocytosed by macrophages, followed by two additional steps: (1) the inflammasome-dependent lysis of infected macrophages to release bacteria to facilitate epithelial invasion (Schnupf and Sansonetti, 2019; Suzuki et al., 2007; Zychlinsky et al., 1994; Zychlinsky et al., 1996), and (2) the concomitant processing and release of IL-1β, a pro-inflammatory cytokine, that drives inflammation (Arondel et al., 1999; Sansonetti et al., 1995; Sansonetti et al., 2000). However, the roles of these particular steps during mammalian oral infection have never been addressed experimentally.

To evaluate the role of NLRC4 inflammasome activation in myeloid cells, we utilized i*Nlrc4Lyz2Cre* mice (Rauch et al., 2017). These mice harbor a germline null mutation in *Nlrc4*, but encode a *Lyz2Cre*-inducible *Nlrc4* cDNA transgene that restores NLRC4 expression selectively in myeloid cells (primarily macrophages, monocytes, and neutrophils). We infected wild-type B6, i*Nlrc4Lyz2Cre*, and B6.*Nlrc4*^*–/–*^ (*Cre*^*–*^) mice and compared disease outcomes across genotypes (Figure 5). Surprisingly, i*Nlrc4Lyz2Cre* mice phenocopied B6.*Nlrc4*^*–/–*^ mice, and did not exhibit significant differences in weight-loss, bacterial colonization of the intestinal epithelium, cecum length, or diarrhea (Figure 5A-E). There was a modest but insignificant increase in inflammatory cytokines CXCL1 and IL-1β in i*Nlrc4Lyz2Cre* mice (Figure 5F, G), but fewer of these mice displayed fecal blood compared to B6.*Nlrc4*^*–/–*^ mice (Figure 5H). These results provide a striking contrast to our previous results with *iNlrc4VilCre* mice in which NLRC4 is selectively expressed in IECs (Mitchell, Roncaioli et al., 2020). Unlike *iNlrc4Lyz2Cre* mice, *iNlrc4VilCre* mice were strongly protected from oral *Shigella* infection, implying that epithelial but not myeloid cell NLRC4 is protective. We conclude that NLRC4-dependent pyroptosis in macrophages is neither a major driver of disease pathogenesis nor bacterial colonization in our oral mouse model of infection.

**Figure 5.**
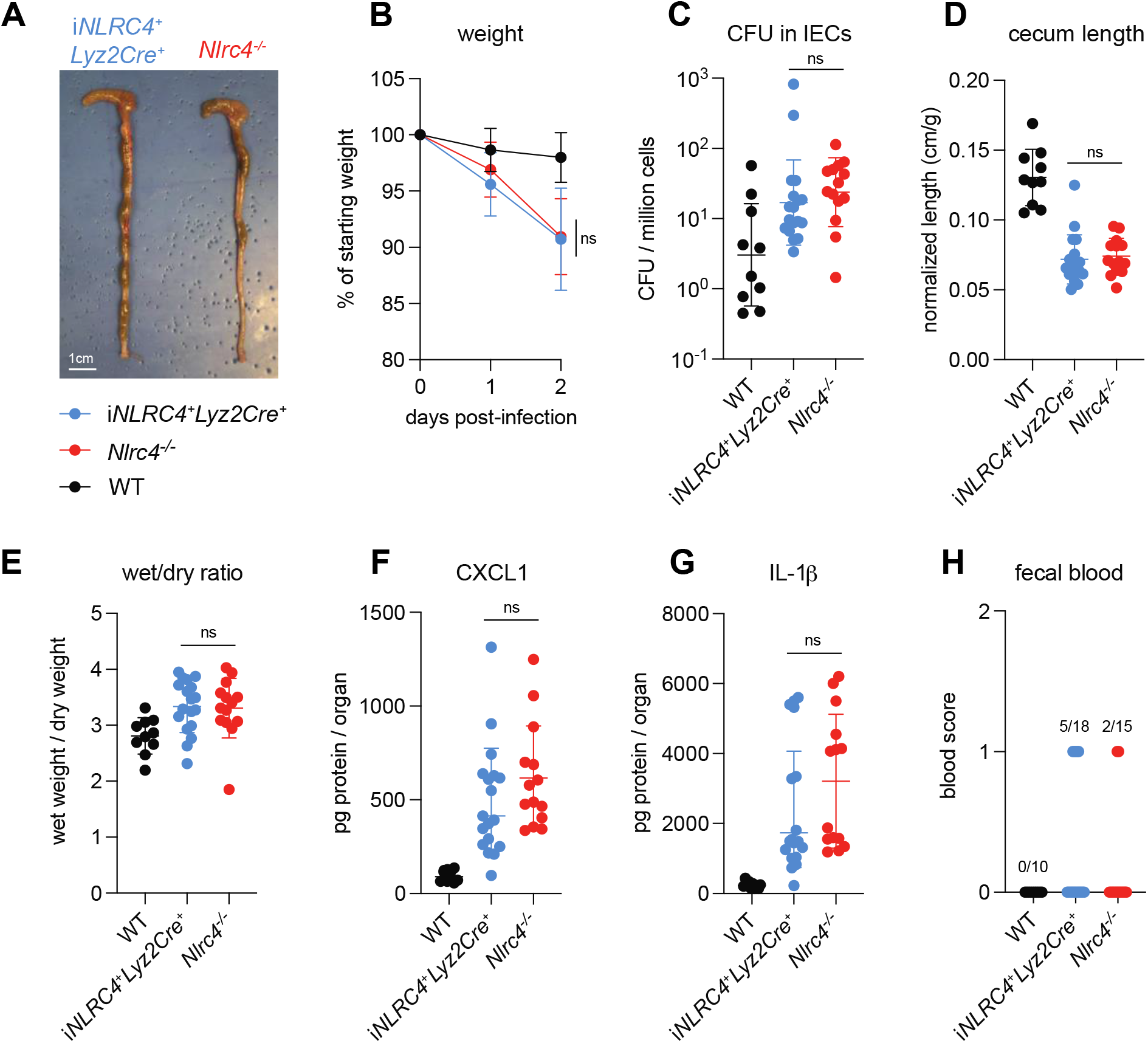
NLRC4 in myeloid-derived cells does not affect *Shigella* pathogenesis. (**A-H**) WT (black) mice and B6.*Nlrc4*^***–/–***^ (Cre^−^) (red) and i*Nlrc4Lyz2Cre* (blue) littermates were treated orally with 25 mg streptomycin sulfate in water and orally challenged the next day with 10^7^ CFU of WT *Shigella flexneri*. Mice were sacrificed at two days post-infection. (**A**) Representative images of the cecum and colon from i*Nlrc4Lyz2Cre* and B6.*Nlrc4*^***–/–***^ mice. Note the similarity in gross pathology between the two genotypes. (**B**) Mouse weights from 0 through 2 days post-infection. Each symbol represents the mean for all mice of the indicated group. (**C**) *Shigella* colony forming units (CFU) per million cells from the combined intestinal epithelial cell (IEC) enriched fraction of gentamicin-treated cecum and colon tissue. (**D**) Quantification of cecum lengths normalized to mouse weight prior to infection; cecum length (cm) / mouse weight (g). (**E**) The ratio of fecal pellet weight when wet (fresh) divided by the fecal pellet weight after overnight drying. Pellets were collected at day two post-infection. (**F, G**) CXCL1 and IL-1β levels measured by ELISA from homogenized cecum and colon tissue of infected mice. (**H**) Blood scores from feces collected at two days post-infection. 1 = occult blood, 2 = macroscopic blood. (**C-H**) Each symbol represents one mouse. Data collected from two independent experiments. Mean ± SD is shown in (**B, D-G**). Geometric mean ± SD is shown in (**C**). Mann-Whitney test, *p < 0.05, **p < 0.01, ***p < 0.001, ****p < 0.0001, ns = not significant (p > 0.05).

IL-1α and IL-1β are related cytokines that are produced downstream of inflammasome activation in myeloid cells and that signal via the common IL-1 receptor. IL-1 cytokines have been implicated in driving inflammation in the context of mouse intranasal *Shigella* challenge (Sansonetti et al., 2000) and rabbit ligated intestinal loop infection (Sansonetti et al., 1995). To better address the role of IL-1 in shigellosis, we crossed B6.*Nlrc4*^*–/–*^ mice to B6.*Il1r1*^*–/–*^ mice to generate B6.*Nlrc4*^*–/–*^*Il1r1*^*–/–*^ double-deficient mice that are susceptible to *Shigella* infection but fail to respond to IL-1. We infected *Nlrc4*^*+/–*^*Il1r1*^*+/–*^, *Il1r1*^*–/–*^, *Nlrc4*^*–/–*^, and *Nlrc4*^*–/–*^*Il1r1*^*–/–*^ littermates and again assessed disease outcomes (Figure 6). Surprisingly, *Nlrc4*^*–/–*^*Il1r1*^*–/–*^ mice phenocopied *Nlrc4*^*–/–*^ mice, and we did not observe significant differences in weight-loss, colonization of the intestinal epithelium, normalized cecum lengths, or inflammatory cytokines (Figure 6A-E). In many bacterial infections, IL-1 signaling initiates the recruitment of neutrophils to sites of infection. However, we did not observe a significant difference in the amount of the neutrophil marker myeloperoxidase (MPO) in the feces of *Nlrc4*^*–/–*^ *Il1r1*^*–/–*^ versus *Nlrc4*^*–/–*^ mice, suggesting that IL-1 might not be essential for neutrophilic inflammation during *Shigella* infection (Figure 6F). We also found that *Nlrc4*-sufficient *Il1r1*^*–/–*^ mice phenocopy WT mice and are resistant to infection. Overall, these results indicate that, despite the increases in IL-1β consistently seen in susceptible mice, IL-1 signaling is not a primary driver of pathogenesis or protection during oral *Shigella* infection. NLRC4-dependent resistance to shigellosis is therefore likely due to the initiation of pyroptosis and expulsion in IECs and not myeloid cell pyroptosis nor IL-1 signaling. Our results leave open a possible role for another inflammasome-dependent cytokine, IL-18, which unlike IL-1β, is highly expressed in IECs.

**Figure 6.**
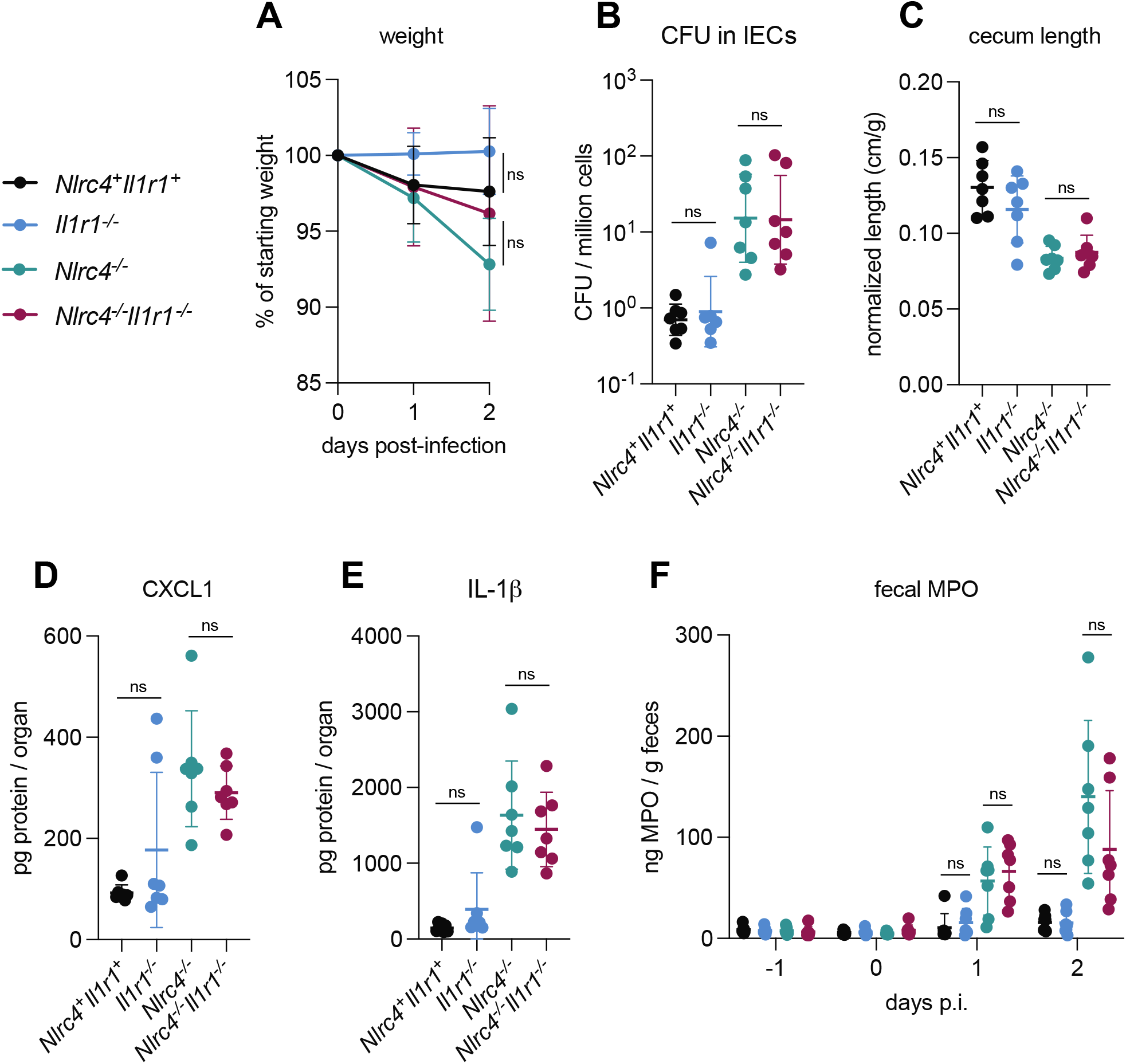
IL-1 signaling does not affect *Shigella* pathogenesis. (**A-F**) WT or *Nlrc4*^*+/–*^*Il1r1*^*+/–*^ (black), *Il1r1*^***–/–***^ (blue), *Nlrc4*^***–/–***^ (teal), and *Nlrc4*^***–/–***^ *Il1r1*^***–/–***^ (maroon) littermates were treated orally with 25 mg streptomycin sulfate in water and orally challenged the next day with 10^7^ CFU of WT *Shigella flexneri*. Mice were sacrificed at two days post-infection. (**A**) Mouse weights from 0 through 2 days post-infection. Each symbol represents the mean for all mice of the indicated group. (**B**) *Shigella* colony forming units (CFU) per million cells from the combined intestinal epithelial cell (IEC) enriched fraction of gentamicin-treated cecum and colon tissue. (**C**) Quantification of cecum lengths normalized to mouse weight prior to infection; cecum length (cm) / mouse weight (g). (**D, E**) CXCL1 and IL-1β levels measured by ELISA from homogenized cecum and colon tissue of infected mice. (**F**) Myeloperoxidase enzyme levels in mouse feces collected each day prior to and during infection and measured by ELISA. (**B-F**) Each symbol represents one mouse. Data are representative of two independent experiments. Mean ± SD is shown in (**A, C-F**). Geometric mean ± SD is shown in (**B**). Mann-Whitney test, *p < 0.05, **p < 0.01, ***p < 0.001, ****p < 0.0001, ns = not significant (p > 0.05).

### TNFα contributes to resistance to *Shigella*

Given that both NLRC4 and CASP11 protect the mouse epithelium from *Shigella* colonization, we reasoned that additional mechanisms of cell death might function in this niche to counteract *Shigella* invasion and spread. Another cell death initiator in the intestine is TNFα, which has been shown to promote *Salmonella*-induced IEC death and dislodgement (Fattinger et al., 2021). TNFα initiates Caspase-8 dependent apoptosis through TNFRI engagement particularly when NF-κB signaling is altered or blocked (Leppkes et al., 2014; Liu et al., 2004; Piguet et al., 1998; Ruder et al., 2019). *Shigella* encodes several effectors reported to inhibit NF-κB signaling (Ashida et al., 2010; Ashida et al., 2013; de Jong et al., 2016; Kim et al., 2005; Newton et al., 2010; Sanada et al., 2012; Wang et al., 2013), and thus, we hypothesized that TNFα might restrict *Shigella* by inducing death of infected IECs.

To assess the *in vivo* role of TNFα during shigellosis, we first infected B6.*Nlrc4*^*–/–*^ mice treated with an antibody that neutralizes TNFα or with an isotype control antibody (Figure 7). B6.*Nlrc4*^*–/–*^ mice that underwent TNFα neutralization were slightly more susceptible to shigellosis than B6.*Nlrc4*^*–/–*^ mice treated with control antibody and displayed modest increases in weight-loss, bacterial burdens in IECs, inflammatory cytokines, and fecal blood (Figure 7A-E). B6.*Nlrc4*^*–/–*^ mice express a functional Caspase-11 inflammasome and given the redundancy we observed between NLRC4 and Caspase-11 (Figures 1-4, (Mitchell, Roncaioli et al., 2020)), we hypothesized that a protective role for TNFα during *Shigella* infection might be most evident in the absence of both of these cell death pathways. To test this, we repeated the experiment in B6.*Nlrc4*^*–/–*^*Casp11*^*–/–*^ mice and, indeed, found that TNFα neutralization on this genetic background significantly increased susceptibility to *Shigella* infection. Mice treated with antibody to TNFα experienced a ∼5% increase in weight-loss, a 10-fold increase in bacterial colonization of the intestinal epithelium, and increases in cecal and colonic shrinkage, diarrhea, inflammatory cytokines, and fecal blood (Figure 7F-O). TNFα levels were elevated significantly in B6.*Nlrc4*^*–/–*^*Casp11*^*–/–*^ mice, indicating that expression of this cytokine is induced in susceptible mice (Figure 7N). The anti-TNFα antibody did not decrease the levels of TNFα measured by ELISA because the antibody neutralizes signaling by the cytokine without interfering with its ability to be detected by ELISA.

**Figure 7.**
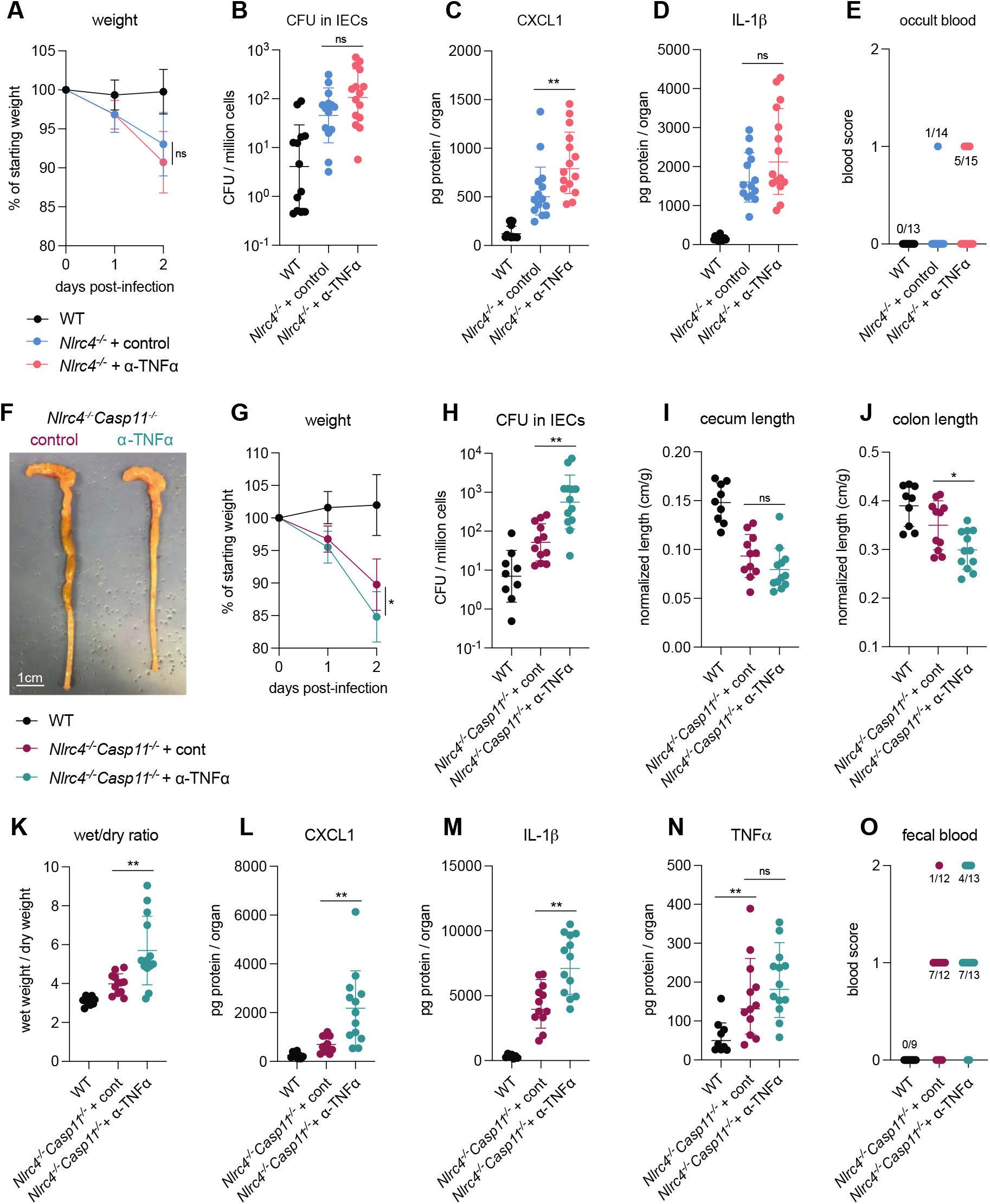
TNFα contributes to resistance to *Shigella* when mice lack NLRC4 and CASP11. B6.WT (black), B6.*Nlrc4*^***–/–***^, and B6.*Nlrc4*^***–/–***^*Casp11*^***–/–***^ mice were treated orally with 25 mg streptomycin sulfate in water and orally challenged the next day with 10^7^ CFU of WT *Shigella flexneri*. In (**A-E**), B6.*Nlrc4*^***–/–***^ mice received 200 μg of either TNFα neutralizing antibody (pink) or isotype control antibody (light blue) by intraperitoneal injection daily from one day before infection through sacrifice at two days post-infection. In (**F-O**), B6.*Nlrc4*^***–/–***^*Casp11*^***–/–***^ mice received 200 μg of either TNFα neutralizing antibody (teal) or isotype control antibody (maroon) by intraperitoneal injection daily from one day before infection through sacrifice at two days post-infection. (**A, G**) Mouse weights from 0 through 2 days post-infection. Each symbol represents the mean for all mice of the indicated group. (**B, H**) *Shigella* colony forming units (CFU) per million cells from the combined intestinal epithelial cell (IEC) enriched fraction of gentamicin-treated cecum and colon tissue. (**C, D, L-N**) CXCL1, IL-1β, and TNFα levels measured by ELISA from homogenized cecum and colon tissue of infected mice. (**E, O**) Blood scores from feces collected at two days post-infection. 1 = occult blood, 2 = macroscopic blood. (**F**) Representative images of the cecum and colon from B6.*Nlrc4*^***–/–***^*Casp11*^***–/–***^ mice receiving either isotype control or TNFα neutralizing antibody. (**I, J**) Quantification of cecum and colon lengths normalized to mouse weight prior to infection; cecum or colon length (cm) / mouse weight (g). (**K**) The ratio of fecal pellet weight when wet (fresh) divided by the fecal pellet weight after overnight drying. Pellets were collected at day two post-infection. (**B-E, H-O**) Each symbol represents one mouse. Data collected from three independent experiments (**A-E**) and two independent experiments (**F-O**). Mean ± SD is shown in (**A, C, D, G, I-N**). Geometric mean ± SD is shown in (**B, H**). Mann-Whitney test, *p < 0.05, **p < 0.01, ***p < 0.001, ****p < 0.0001, ns = not significant (p > 0.05).

Importantly, we could also observe a significant protective role for TNFα in similar experiments performed in 129.*Nlrc4*^*–/–*^ mice that are naturally deficient in Caspase-11 (Figure 7 — figure supplement 1), confirming that TNFα-dependent protection is redundant with both NLRC4 and Caspase-11. These results suggest that a hierarchy of cell death pathways protect the intestinal epithelium from *Shigella* infection. NLRC4 appears to be both necessary and sufficient to protect mice from disease, but in the absence of NLRC4, functional CASP11 can provide compensatory protection. However, in the absence of both NLRC4 and Caspase-11, a critical role for TNFα is revealed (Figure 2 — figure supplement 2).

### Loss of multiple cell death pathways renders mice hyper-susceptible to *Shigella*

To directly test the role of Caspase-8-dependent cell death during *Shigella* infection, we generated mice lacking either Caspases-1 and 11 (B6.*Casp1/11*^*–/–*^*Ripk3*^*–/–*^), Caspase-8 (B6.*Casp8*^*–/–*^*Ripk3*^*–/–*^), or Caspases-1, 11, and 8 (B6.*Casp1/11/8*^*–/–*^*Ripk3*^*–/–*^). Since loss of Caspase-8 results in Ripk3-depedent embryonic lethality, all three genotypes also lack *Ripk3. Casp1/11*^*–/–*^*Ripk3*^*–/–*^ mice retain Caspase-8 function downstream of both NLRC4 and TNFα (Figure 2 — figure supplement 2) and based on our previous experiments with *Casp1/11*^−/–^ mice (Mitchell, Roncaioli et al., 2020), we expected that these mice would be resistant to infection. Similarly, *Casp8*^*–/–*^*Ripk3*^*–/–*^ mice retain the ability to recruit Caspase-1 to NLRC4 and to initiate cell death via Caspase-11 ((Figure 2 — figure supplement 2) and should also thus be resistant to infection. *Casp1/11/8*^*–/–*^*Ripk3*^*–/–*^ mice, however, should lack the cell death pathways initiated by NLRC4 (via Caspase-1 or Caspase-8), Caspase-11, and TNFα (Figure 2 — figure supplement 2), and our results above suggest that these mice might be highly susceptible to infection.

We infected wild-type B6 mice as well as *Casp1/11*^*–/–*^*Ripk3*^*–/–*^, *Casp8*^*–/–*^*Ripk3*^*–/–*^, and *Casp1/11/8*^*–/–*^ *Ripk3*^*–/–*^ littermates and assessed disease phenotypes across all four genotypes (Figure 8). We found that *Casp8*^*– /–*^*Ripk3*^*–/–*^ mice largely phenocopied wild-type B6 mice, and were resistant to infection, exhibiting minimal weight-loss, diarrhea, cecal or colonic shrinkage, and fecal blood (Figure 8A, B, D, E, F, I). Furthermore, we could not detect significant increases in bacterial burdens in the epithelium (Figure 8C) nor inflammatory cytokines (Figure 8G, H) in *Casp8*^*–/–*^*Ripk3*^*–/–*^ mice. These results suggest that Caspase-8 alone is not necessary for resistance to *Shigella* in the presence of functional NLRC4–CASP1 and CASP11 inflammasomes.

**Figure 8.**
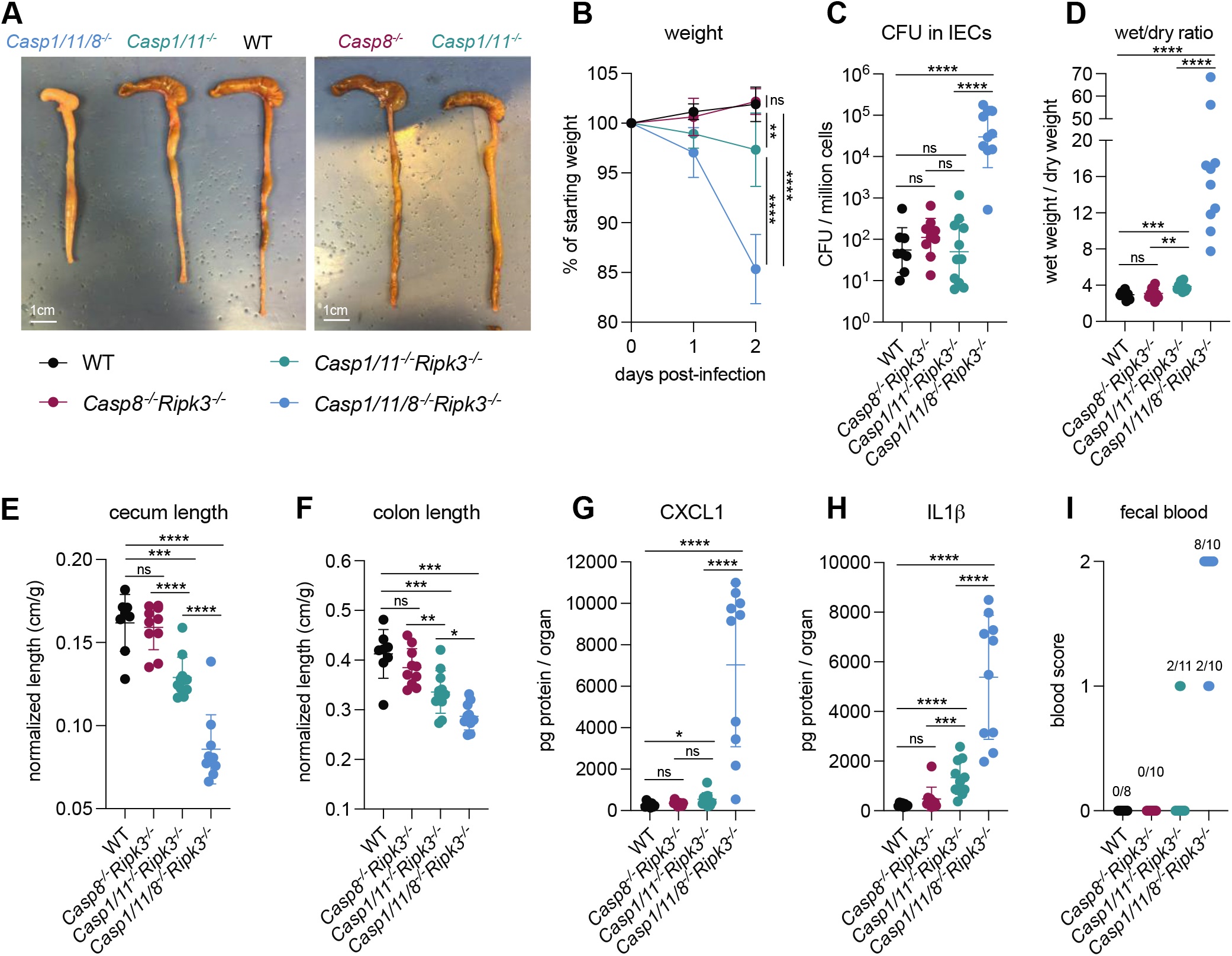
Loss of multiple cell death pathways renders mice hyper-susceptible to *Shigella*. (**A-I**) B6.WT mice (black), and B6.*Casp8*^***–/–***^*Ripk3*^***–/–***^ (maroon), B6.*Casp1/11*^***–/–***^*Ripk3*^***–/–***^ (teal), and B6.*Casp1/11/8*^***–/–***^*Ripk3*^***–/–***^ (light blue) littermates were treated orally with 25 mg streptomycin sulfate in water and orally challenged the next day with 10^7^ CFU of WT *Shigella flexneri*. Mice were sacrificed at two days post-infection. (**A**) Representative images of the cecum and colon of infected B6.WT, B6.*Casp8*^***–/–***^*Ripk3*^***–/–***^, B6.*Casp1/11*^***–/–***^*Ripk3*^***–/–***^, and *Casp1/11/8*^***–/–***^*Ripk3*^***–/–***^ mice. Note the severe inflammation in the *Casp1/11/8*^***–/–***^ *Ripk3*^***–/–***^ mice (left-most organs). (**B**) Mouse weights from 0 through 2 days post-infection. Each symbol represents the mean for all mice of the indicated group. (**C**) *Shigella* colony forming units (CFU) per million cells from the combined intestinal epithelial cell (IEC) enriched fraction of gentamicin-treated cecum and colon tissue. (**D**) The ratio of fecal pellet weight when wet (fresh) divided by the fecal pellet weight after overnight drying. Pellets were collected at day two post-infection. (**E, F**) Quantification of cecum and colon lengths normalized to mouse weight prior to infection; cecum or colon length (cm) / mouse weight (g). (**G, H**) CXCL1 and IL-1β levels measured by ELISA from homogenized cecum and colon tissue of infected mice. (**I**) Blood scores from feces collected at two days post-infection. 1 = occult blood, 2 = macroscopic blood. (**C-I**) Each symbol represents one mouse. Data collected from two independent experiments. Mean ± SD is shown in (**B, D-H**). Geometric mean ± SD is shown in (**C**). Mann-Whitney test, *p < 0.05, **p < 0.01, ***p < 0.001, ****p < 0.0001, ns = not significant (p > 0.05).

Interestingly, *Casp1/11*^*–/–*^*Ripk3*^*–/–*^ mice were not fully resistant to disease and experienced modest weight-loss (∼5% relative to WT), diarrhea, cecal and colonic shrinkage, and a small but significant increase in inflammatory cytokines CXCL1 and IL-1β (Figure 8A, B, D-H). Two of the eleven *Casp1/11*^*–/–*^*Ripk3*^*–/–*^ mice also exhibited occult fecal blood (Figure 8I). Despite the modest susceptibility of this genotype, we were unable to detect an increase in bacterial colonization of the intestinal epithelium relative to WT or *Casp8*^*–/–*^*Ripk3*^*–/–*^ (Figure 8C).

The most striking observation was that *Casp1/11/8*^*–/–*^*Ripk3*^*–/–*^ mice were highly susceptible to *Shigella* infection, exhibiting severe weight-loss (∼15% of starting weight), diarrhea, and cecal and colonic shrinkage (Figure 8A, B, D-F). These mice also exhibited a massive (>500×) increase in bacterial colonization of the epithelium (Figure 8C) and elevated levels of inflammatory cytokines (Figure 8G, H). All *Casp1/11/8*^*–/–*^*Ripk3*^*–/–*^ mice presented with blood in their feces (Figure 8I) and one of the ten mice also died of shigellosis within 2 days of infection. The ceca and colons of *Casp1/11/8*^*–/–*^*Ripk3*^*–/–*^ mice were highly inflamed – the tissue thickened, turned white, and sections of the epithelium appeared to have been shed into the lumen, which was completely devoid of feces and filled instead with neutrophilic pus (Figure 8A). While the most significant inflammation in B6.*Nlrc4*^*–/–*^ mice is typically seen in the cecum (Mitchell, Roncaioli et al., 2020), we noted that the colon of *Casp1/11/8*^*–/–*^*Ripk3*^*–/–*^ mice was highly inflamed as well (Figure 8A, F), suggesting that a protective role for Caspase-8 might be most important in this organ. Taken together, our results imply that redundant cell death pathways protect mice from disease upon oral *Shigella* challenge. Genetic removal of three caspases essential to this response leads to severe disease and even death. However, removal of one or two caspases critical to this response does not lead to severe disease because of significant compensation from the other pathway(s). We observe a hierarchical importance of the cell death pathways, namely, NLRC4 > CASP11 > TNFα–CASP8 (Figure 2 — figure supplement 2). We speculate that this hierarchy may be established by the timing by which a pathway can sense invasive *Shigella* within the epithelium.

## Discussion

We have previously shown that intestinal epithelial cell expression of the NAIP–NLRC4 inflammasome is sufficient to confer resistance to shigellosis in mice (Mitchell, Roncaioli et al., 2020). Activation of NAIP– NLRC4 by *Shigella* drives pyroptosis and expulsion of infected IECs. Genetic removal of NAIP–NLRC4 from IECs allows *Shigella* to colonize the intestinal epithelium, an event which drives intestinal inflammation and disease. Mouse IECs, however, deploy additional initiators of programmed cell death (Patankar and Becker, 2020) and it remained an open question whether these cell death pathways might also counteract *Shigella*.

We utilized the natural variation in 129.*Nlrc4*^*–/–*^ mice, which lack functional CASP11 (Kayagaki et al., 2011), to show that CASP11 partially controls the difference in susceptibility between 129.*Nlrc4*^*–/–*^ and B6.*Nlrc4*^*–/–*^ mice (Figure 1, Figure 1 — figure supplement 1). In F_1_ 129/B6.*Nlrc4*^*–/–*^ × 129.*Nlrc4*^*–/–*^ backcrossed mice, which were either 129/129 or B6/129 at the *Casp11* locus, increased disease severity and colonization of the intestinal epithelium was associated with a homozygous null *Casp11*^129^ locus. We also investigated the role of *Hiccs*, a locus present in 129 mice that confers increased susceptibility to *Helicobacter hepaticus*-induced colitis (Boulard et al., 2012). The 129 *Hiccs* locus contains polymorphisms in the *Alpk1* gene which encodes alpha-kinase 1 (ALPK1), an activator of NF-κB which has been show to sense *Shigella-*derived ADP-heptose in human cells (Zhou et al., 2018). However, we did not find that *Hiccs* contributed to differences in susceptibility between the two strains (Figure 1 — figure supplement 2).

We observed that *ΔospC3 Shigella* is significantly attenuated in B6.*Nlrc4*^*–/–*^ mice but not in B6.*Nlrc4*^*–/–*^ *Casp11*^*–/–*^, indicating by a “genetics squared” analysis (Persson and Vance, 2007) that *Shigella* effector OspC3 inhibits CASP11 during oral mouse infection (Figures 3, 4). The striking decrease in colonization of the intestinal epithelium in *ΔospC3*-infected B6.*Nlrc4*^*–/–*^ mice relative to *ΔospC3*-infected B6.*Nlrc4*^*–/–*^*Casp11*^*–/–*^ mice suggests that CASP11-dependent protection is epithelial intrinsic. *Shigella* also deploys an effector, IpaH7.8, which degrades human (but not mouse) GSDMD to block pyroptosis, further underscoring the importance of this axis in defense (Luchetti et al., 2021). We note that CASP11-dependent protection is not sufficient to render *ΔospC3*-infected B6.*Nlrc4*^*–/–*^ mice completely resistant to disease symptoms, perhaps because the priming required to induce CASP11 expression might delay its protective response (Oh et al., 2021).

Despite its role as a key cell death initiator in the gut (Patankar and Becker, 2020; Piguet et al., 1998; Ruder et al., 2019), TNFα has not yet been shown to play a major role in defense against pathogens that colonize the intestinal epithelium. Indeed, its role is usually detrimental to the host. For example, TNFα is a major driver of pathology during Crohn’s Disease (van Dullemen et al., 1995). In the context of *Salmonella* infection, TNFα appears to drive widespread pathological death and dislodgement of IECs at 72 hours post-infection (Fattinger et al., 2021). Here, we show that TNFα is protective during oral *Shigella* infection, providing a rationale for why this cytokine is produced in the intestine. In both B6.*Nlrc4*^*–/–*^*Casp11*^*–/–*^ and 129.*Nlrc4*^*–/–*^ mice, TNFα neutralization lead to an increase in severity of infection and a 10-fold increase in bacterial colonization of the intestinal epithelium, suggesting that TNFα-dependent IEC apoptosis restricts colonization of this niche by *Shigella* (Figure 7). NF-κB-dependent cytokines IL-1β and CXCL1 increase after TNFα neutralization, indicating that protection is not likely driven by the TNFα-dependent activation of NF-κB. An important next step will be to assess whether there is a link between the *Shigella-*dependent inhibition of NF-κB (Ashida et al., 2010; Ashida et al., 2013; de Jong et al., 2016; Kim et al., 2005; Newton et al., 2010; Sanada et al., 2012; Wang et al., 2013) and CASP8-dependent apoptosis in IECs. Interestingly, TNFα neutralization in B6.*Nlrc4*^−/–^ mice had only a modest effect on disease susceptibility and colonization of IECs, indicating that TNFα-dependent protection is only fully revealed in the absence of both NLRC4 and CASP11. The protective cell death pathway hierarchy (NLRC4 > CASP11 > TNFα–CASP8) established by these experiments highlights the importance of redundant layers of immunity as a strategy to counteract pathogen evolution.

We find that *Casp1/11/8*^*–/–*^*Ripk3*^*–/–*^ mice, which lack the pathways to execute pyroptosis, extrinsic apoptosis, and necroptosis, experience severe *Shigellosis* with a 500-fold increase in colonization of the intestinal epithelium relative to B6 wild-type mice (Figure 8). Although we did not directly compare the two mouse strains, *Casp1/11/8*^*–/–*^*Ripk3*^*–/–*^ mice (Figure 8) experienced more severe disease and epithelial colonization than *Nlrc4*^*–/–*^*Casp11*^*–/–*^ mice (Figure 2, 4, 7). We speculate that the additional susceptibility of *Casp1/11/8*^*–/–*^*Ripk3*^*–/–*^ mice partially results from the absence of TNFRI–CASP8-dependent apoptosis and possibly from the absence of RIPK3-dependent necroptosis. However, there likely exist other roles for CASP8 during *Shigella* infection which might further account for its importance (Gitlin et al., 2020; Philip et al., 2016; Schwarzer et al., 2020; Stolzer et al., 2022; Weng et al., 2014; Woznicki et al., 2021). Interestingly, *Casp1/11*^*–/–*^ *Ripk3*^*–/–*^ mice experience modest susceptibility to *Shigella* relative to *Casp8*^*–/–*^*Ripk3*^*–/–*^ mice, which are fully protected (Figure 8), potentially because NLRC4–CASP8-dependent cell death is delayed relative to NLRC4– CASP1-dependent cell death (Lee et al., 2018; Rauch et al., 2017). Furthermore, CASP8 might be both protective and partially inhibited by a *Shigella* effector (Ashida et al., 2020), thus implicating RIPK3-dependent IEC necroptosis in protection (Wen et al., 2017). In the future, a comparison of *Casp1/11*^*–/–*^ and *Casp1/11*^*–/–*^ *Ripk3*^*–/–*^ mice could test whether RIPK3-dependent necroptosis provides an additional layer of protection during *Shigella* infection.

Despite the commonly held belief that macrophage pyroptosis and IL-1β release drive *Shigella* pathogenesis (Schnupf and Sansonetti, 2019; Schroeder and Hilbi, 2008), we find no major protective or pathogenic role for either during *Shigella* infection (Figure 5, 6). These data suggest that epithelial-specific cell death and expulsion may be the key mechanism that protects mice from *Shigella*. Infections in IL-18 deficient mice will further clarify the role of inflammasome-dependent cytokines in protection. Additional studies in bone marrow chimeric mice or tissue specific knockout mice are required to genetically confirm whether the protective effects of CASP11 and TNFα are epithelial intrinsic.

Here, we illustrate the existence of a layered cell death pathway hierarchy that is essential in defense against oral *Shigella* infection in mice. Our work highlights the significant evolutionary steps required by *Shigella* to overcome these pathways and cause disease in humans. We observed a correlation between bacterial burdens in IECs and pathogenicity in our experiments, indicating that the extent to which *Shigella* can colonize the intestinal epithelium dictates the severity of disease during infection. However, the sensors within IECs that initiate inflammation and drive pathogenicity *in vivo* have yet to be uncovered and might present an ideal pharmacological target to limit pathological inflammation during acute *Shigella* infection.

## Materials and Methods

### Key resources table

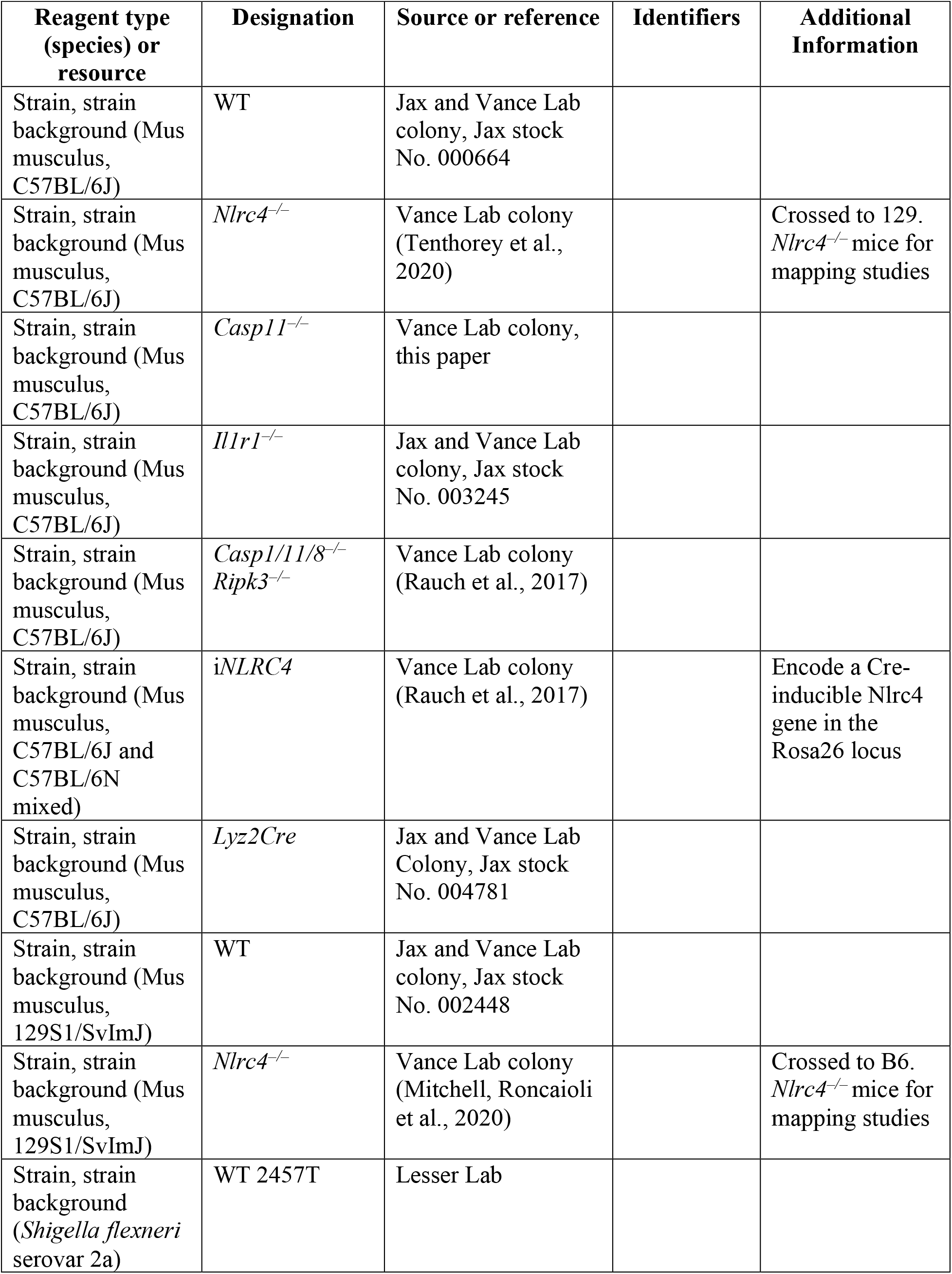
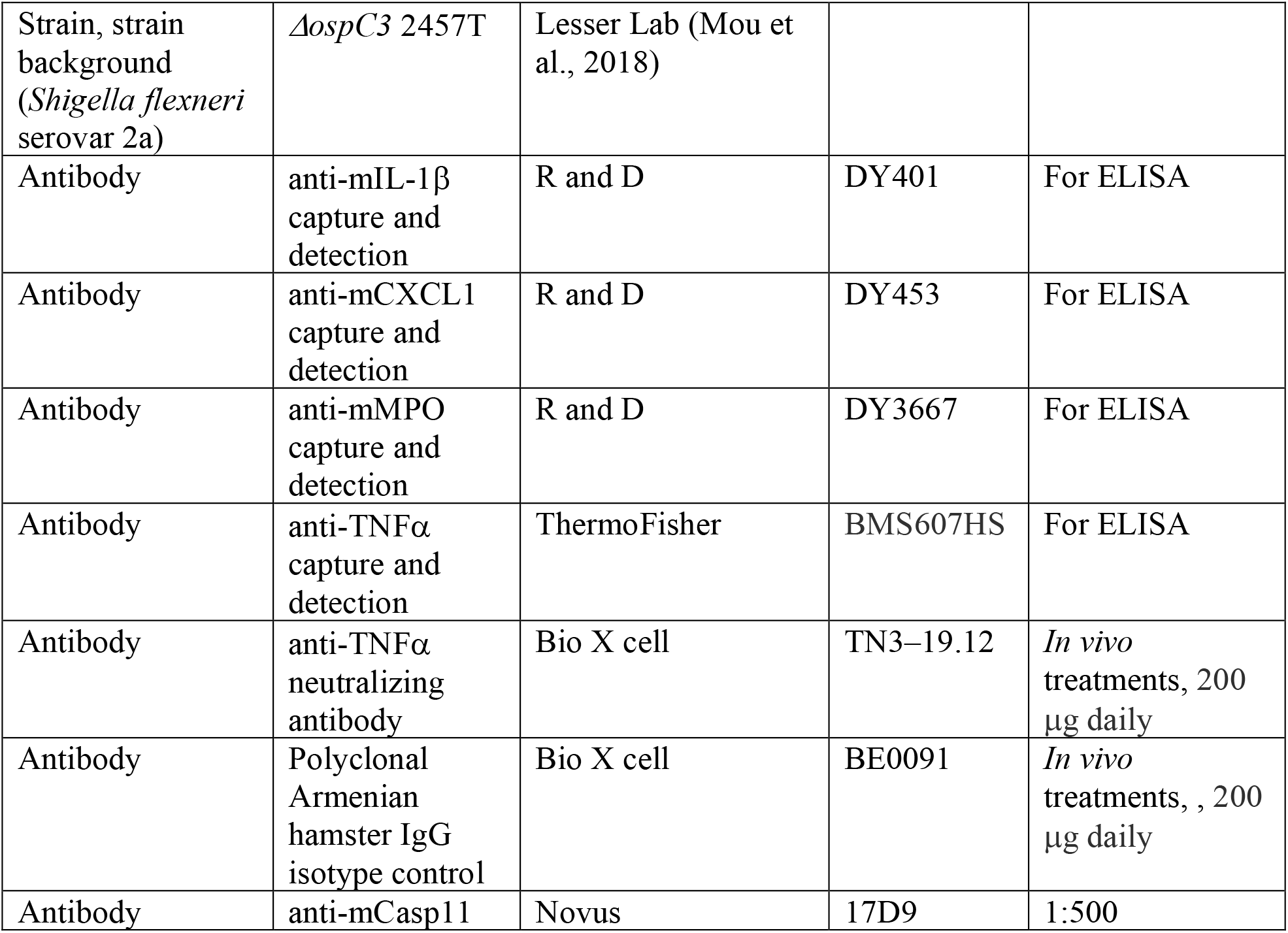

### Animal Procedures

All mice were maintained in a specific pathogen free colony until 1–8 weeks prior to infection, maintained under a 12 hr light-dark cycle (7 am to 7 pm), and given a standard chow diet (Harlan irradiated laboratory animal diet) ad libitum. Animals used in infection experiments were littermates or, if not possible, were generally cohoused upon weaning. In cases when mice were not co-housed upon weaning, mice were cohoused for at least three week prior to infection. Different experimental treatments (comparing disease across different *Shigella* genotypes or antibody treatments) were stratified within mouse genotypes of the same litter, where possible, to ensure that phenotypes were not the result of the differences in different litter microbiomes. Mice were transferred from a SPF colony to an ABSL2 facility at least one week prior to infection. All mouse infections complied with the regulatory standards of, and were approved by, the University of California, Berkeley Animal Care and Use Committee. B6.*Nlrc4*^*–/–*^ and 129.*Nlrc4*^*–/–*^ mice were generated as previously described (Mitchell, Roncaioli et al., 2020; Tenthorey et al., 2020). F_1_ 129/B6.*Nlrc4*^*–/–*^ were generated by crossing parental 129.*Nlrc4*^*–/–*^ and B6.*Nlrc4*^*–/–*^ mice. F_1_ 129/B6.*Nlrc4*^*–/–*^ mice were crossed to parental 129.*Nlrc4*^*–/–*^ mice to generate backcrossed mice that were either B6/129 or 129/129 at all loci. 129 and B6 *Casp11* alleles were distinguished by PCR and sequencing using the primers B6.129_Casp11_F 5’ GTTATCTATCAGTAGGAAGTGG 3’ and B6.129_Casp11_R 5’ AAACTAATACTTCTTATGAGAGC 3’. 129 mice have a distinguishable 5-bp deletion encompassing the exon 7 splice acceptor junction (Kayagaki et al., 2011). The *Hiccs* locus was genotyped by PCR using the primers D3Mit348_F 5’ CATCATGCATACTTTTTTCCTCA 3’, D3Mit348_R 5’ GCCAAATCATTCACAGCAGA 3’, D3Mit319_F 5’ TCTCCCTCACTTTTTCCTTCC 3’, and D3Mit319_R 5’ AACAGCCAGTCCAGCAAATC 3’ to distinguish polymorphisms between the B6 and 129 alleles. B6.*Nlrc4*^*–/–*^*Casp11*^*–/–*^ animals were generated by targeting *Casp11* via CRISPR-Cas9 mutagenesis in existing B6.*Nlrc4*^*–/–*^ mice. CRISPR/Cas9 targeting was performed by electroporation of Cas9 protein and sgRNA into fertilized zygotes, essentially as described previously (Chen et al., 2016). Founder mice were genotyped by PCR and sequencing using the primers: Casp4_F 5’ GTCTTTAGCCCTTGAGAAGGACAC 3’ and Casp4_R 5’ CACCCCTTCACTTGAGTTTCTCC 3’. Founders carrying mutations were bred one generation to B6.*Nlrc4*^*–/–*^ mice to separate modified haplotypes. Homozygous lines were generated by interbreeding heterozygotes carrying matched haplotypes. i*Nlrc4* mice (Rauch et al., 2017) were previously described. i*Nlrc4* mice were crossed to the B6.*Nlrc4*^*–/–*^ line and then further crossed to *Lyz2Cre* (Jax strain 004781) transgenic lines on a B6.*Nlrc4*^*–/–*^ background to generate i*Nlrc4Lyz2Cre* mice. *Nlrc4*^*–/–*^*Il1r1*^*–/–*^ mice were generated by crossing B6.*Nlrc4*^*–/–*^ mice to B6.*Il1r1*^*–/–*^ mice (Jax strain 003245). B6.*Casp8*^*–/–*^*Ripk3*^*–/–*^, B6.*Casp1/11*^*–/–*^*Ripk3*^*–/–*^, and B6.*Casp1/11/8*^*–/–*^*Ripk3*^*–/–*^ mice were generated as previously described (Rauch et al., 2017).

### *Shigella* Strains

Mouse infections were conducted with the *Shigella flexneri* serovar 2a 2457T strain, WT or *ΔospC3* (Mou et al., 2018). Natural streptomycin resistant strains of WT and *ΔospC3* were generated by plating cultured bacteria on tryptic soy broth (TSB) plates containing 0.01% Congo Red (CR) and increasing concentrations of streptomycin sulfate. Streptomycin-resistant strains were confirmed to grow indistinguishably from parental strains in TSB broth lacking antibiotics, indicating an absence of streptomycin-dependence.

### *In Vivo Shigella* Infections and Treatments

Streptomycin resistant *Shigella flexneri* was grown at 37°C on tryptic soy agar plates containing 0.01% Congo red (CR), supplemented with 100 μg/mL of streptomycin sulfate. For infections, a single CR-positive colony was inoculated into 5 mL TSB and grown shaking overnight at 37°C. Saturated cultures were back-diluted 1:100 in 5 mL fresh TSB shaking for 2-3 hours at 37°C. The approximate infectious dose was determined by spectrophotometry (OD_600_ of 1 = 10^8^ CFU/mL). Bacteria were pelleted at 5000×*g*, washed twice in PBS, and suspended in PBS for infection by oral gavage. Actual infectious dose was determined by serially diluting a fraction of the initial inoculum and plating on TSB plates containing 0.01% CR and 100 μg/mL streptomycin. Mouse infections were performed in 6–22 week old mice. Initially, mice deprived of food and water for 4–6 hours were orally gavaged with 100 μL of 250 mg/mL streptomycin sulfate dissolved in water (25 mg/mouse) and placed in a cage with fresh bedding. One day later, mice again deprived of food and water for 4–6 hours were orally gavaged with 100 μL of log-phase, streptomycin resistant *Shigella flexneri* suspended in PBS at a dose of 10^8^ CFU/mL (10^7^ CFU/mouse). Mouse weights and fecal pellets were recorded or collected daily from 1 day prior to infection to the day of euthanasia and harvest to assess the severity of disease and biomarkers of inflammation. Fecal colonization (CFU/gram of feces) and successful challenge were determined by homogenizing feces collected one day post-infection and plating (see below). For *in vivo* antibody treatments, 200 μg of anti-TNFα antibody (Bio X Cell, clone TN3–19.12) and polyclonal Armenian hamster IgG isotype control antibody (Bio X Cell) were administered by intraperitoneal injection daily starting one day prior to infection.

### Fecal CFUs, fecal MPO ELISAs, wet/dry ratio, fecal blood

Fecal pellets were collected in 2 mL tubes, suspended in 1 mL of 2% FBS in PBS containing protease inhibitors, and homogenized using a polytron homogenizer at 18,000 rpm. For CFU enumeration, serial dilutions were made in PBS and plated on TSB plates containing 0.01% CR and 100 mg/mL streptomycin sulfate. For MPO ELISAs, fecal homogenates were spun at 2000×*g* and supernatants were plated in duplicate on absorbent immunoassay 96-well plates. Recombinant mouse MPO standard, MPO capture antibody, and MPO sandwich antibody were purchased from R&D Systems. Wet/dry ratios were determined by weighing fecal pellets before and after they had been dried in a fume hood. The presence or absence of fecal blood in fresh pellets was determined by macroscopic observation or by applying wet fecal samples to detection tabs from a Hemoccult blood testing kit (Beckman Coulter).

### Intestinal CFU determination

To enumerate intracellular *Shigella* CFU from the intestinal epithelial cell fraction of mouse ceca and colons, organs were removed from mice upon sacrifice, cut longitudinally and removed of luminal contents by washing in PBS. Tissues were place in 14 mL culture tubes, incubated in RPMI with 5% FBS, 2 mM L-glutamine, 25 mM HEPES, and 400 μg/mL of gentamicin for 1–2 hours, and vortexed briefly. Tissues were then washed five times in PBS, cut into 1 cm pieces, placed in 15 mL of stripping solution (HBSS, 10 mM HEPES, 1 mM DTT, 2.6 mM EDTA), and incubated at 37°C for 25 min with gentle agitation. Supernatants were passed through a 100 micron filter and the remaining pieces of tissue were shaken vigorously in a 50 mL conical with 10 mL of PBS and passed again through the 100 micron filter. This enriched epithelial cell fraction was incubated in 50 μg/mL gentamicin for 30-40 minutes on ice, spun at 300×*g* at 4°C for 8 min, and washed twice by aspirating the supernatant, resuspending in PBS, and spinning at 300×*g* at 4°C for 5 min. After the first wash, a fraction of cells were set aside to determine the cell count. After the second wash, the pellet was resuspended and lysed in 1 mL of 1% Triton X-100. Serial dilutions were made from this solution and plated on TSB agar plates containing 0.01% CR and 100 μg/ml streptomycin and CR+ positive colonies were counted following overnight incubation at 37°C.

### Tissue ELISAs

After isolating the intestinal epithelial cell fraction (above), the remaining tissue was transferred to a 14 mL culture tube containing 1 mL of PBS containing 2% FBS and protease inhibitors. Organs were homogenized using a polytron homogenizer at 20,000 rpm, centrifuged at 2000×*g*, and supernatants were plated on absorbent immunoassay 96-well plates. Recombinant mouse CXCL1 and IL-1β standards, capture antibodies, and sandwich antibodies were purchased from R and D. TNFα levels were detected using a high sensitivity ELISA from ThermoFisher (order no: BMS607HS).

### Immunoblot and antibodies

Lysates were prepared from *Casp11*^*+/–*^ and *Casp11*^*–/–*^ mouse bone marrow derived macrophages and clarified by spinning at 16,100×*g* for 10 min at 4°C. Clarified lysates were denatured in SDS loading buffer. Samples were separated on NuPAGE Bis-Tris 4–12% gradient gels (ThermoFisher) following the manufacturer’s protocol.

Proteins were transferred onto Immobilon-FL PVDF membranes at 375mA for 90 min and blocked with Odyssey blocking buffer (Li-Cor). Proteins were detected on a Li-Cor Odyssey Blot Imager using an anti-Caspase-11 primary antibody (cone 17D9) and Alexfluor-680 conjugated secondary antibody (Invitrogen).

## Acknowledgements

We thank P Mitchell, I Rauch, S Fattinger, and K Eislmayr for discussion and advice. We are grateful to members of the Vance and Barton Labs for discussions. Funding: REV is an HHMI Investigator and is supported by NIH AI075039 and AI063302. JLR is an Irving H Wiesenfeld CEND Fellow; EAT is supported by the UC Berkeley Department of Molecular and Cell Biology NIH Training Grant 5T32GM007232-42; CFL is a Brit d’Arbeloff MGH Research Scholar and supported by NIH AI064285 and NIH AI128743.

## Additional Information

### Author Contributions

Justin L Roncaioli, conceptualization, formal analysis, validation, investigation, methodology, writing – original draft, writing – review and editing; Janet Peace Babirye, investigation; Roberto A Chavez; investigation, Elizabeth A Turcotte, investigation; Fitty Liu, investigation; Angus Y Lee, resources; Cammie F Lesser, resources, supervision, funding acquisition, methodology, writing – review and editing; Russell E Vance, conceptualization, resources, supervision, funding acquisition, methodology, writing – original draft, writing – review and editing.

### Ethics

Animal experimentation: This study was performed in strict accordance with the recommendations in the Guide for the Care and Use of Laboratory Animals of the National Institutes of Health. All of the animals were handled according to approved institutional animal care and use committee (IACUC) protocols (AUP-2014-09-6665-1) of the University of California Berkeley.

---

**Figure 1 — figure supplement 1.**
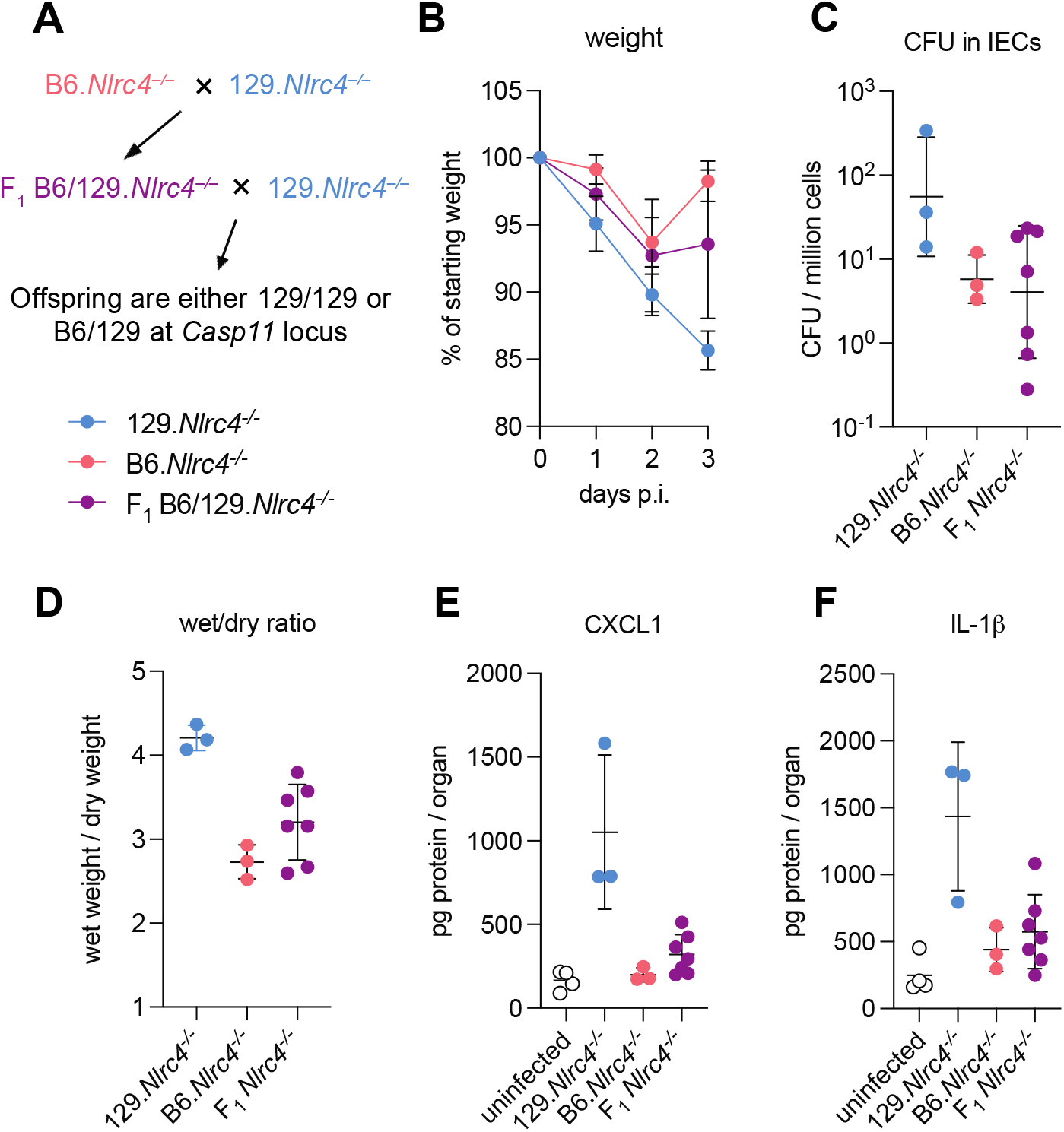
B6/129.*Nlrc4*^*–/–*^ F_1_ hybrids are modestly susceptibility to *Shigella*. Crossing scheme to generate B6/129.*Nlrc4*^*–/–*^ F_1_ mice and backcrossed *Nlrc4*^*–/–*^ mice that are heterozygous B6/129 or homozygous 129/129 at *Casp11*. (**B-E**) B6.*Nlrc4*^***–/–***^ (pink), 129.*Nlrc4*^***–/–***^ (light blue), and 129/B6.*Nlrc4*^***–/–***^ F_1_ (plum) mice were treated orally with 25 mg streptomycin sulfate in water and orally challenged the next day with 10^7^ CFU of WT *Shigella flexneri*. Mice were sacrificed at three days post-infection. (**B**) Mouse weights from 0 through 3 days post-infection. Each symbol represents the mean for all mice of the indicated group. (**C**) *Shigella* colony forming units (CFU) per million cells from the combined intestinal epithelial cell (IEC) enriched fraction of gentamicin-treated cecum and colon tissue. (**D**) The ratio of fecal pellet weight when wet (fresh) divided by the fecal pellet weight after overnight drying. A larger wet/dry ratio indicates increased diarrhea. Pellets were collected at day two post-infection. (**E, F**) CXCL1 and IL-1β levels measured by ELISA from homogenized cecum and colon tissue of infected mice. (**C-F**) Each symbol represents one mouse. Mean ± SD is shown in (**B, D-F**). Geometric mean ± SD is shown in (**C**).

**Figure 1 — figure supplement 2.**
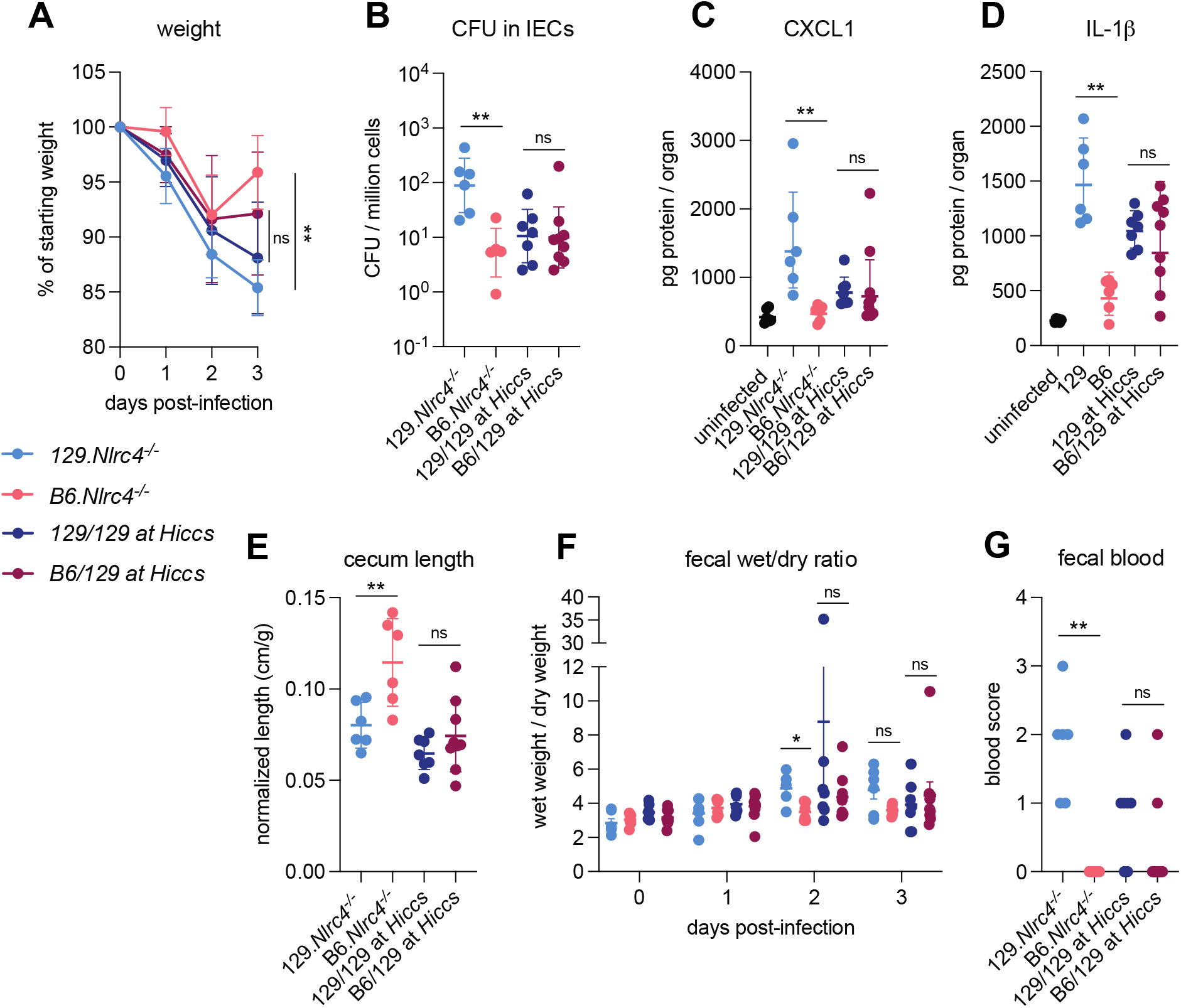
*Hiccs* does not contributes to resistance of B6 versus 129 *Nlrc4*^*–/–*^ mice to shigellosis. (**A-G**) B6.*Nlrc4*^*–/–*^ (pink), 129.*Nlrc4*^*–/–*^ (light blue), backcrossed littermates that are homozygous 129/129 at *Hiccs* (dark blue), and backcrossed littermates that are heterozygous B6/129 at *Hiccs* (maroon) were co-housed for 3 weeks, treated orally with 25 mg streptomycin sulfate in water, and orally challenged the next day with 10^7^ CFU of WT *Shigella flexneri*. Mice were sacrificed at three days post-infection. (**A**) Mouse weights from 0 through 3 days post-infection. Each symbol represents the mean for all mice of the indicated genotype. (**B**) *Shigella* colony forming units (CFU) per million cells from the combined intestinal epithelial cell (IEC) enriched fraction of gentamicin-treated cecum and colon tissue. (**C, D**) CXCL1 and IL-1β levels measured by ELISA from homogenized cecum and colon tissue of infected mice. (**E**) Quantification of cecum lengths normalized to mouse weight prior to infection; cecum length (cm) / mouse weight (g). (**F**) The ratio of fecal pellet weight when wet (fresh) divided by the fecal pellet weight after overnight drying. A larger wet/dry ratio indicates increased diarrhea. Pellets were collected daily from 0-3 days post-infection. (**G**) Additive blood scores from feces collected at two and three days post-infection. 1 = occult blood, 2 = macroscopic blood for a given day. (**B-G**) Each symbol represents one mouse. Mean ± SD is shown in (**A, C-E**). Geometric mean ± SD is shown in (**B**). Mean ± SEM is show in (**F**). Mann-Whitney test, *p < 0.05, **p < 0.01, ***p < 0.001, ****p < 0.0001, ns = not significant (p > 0.05)

**Figure 2 — figure supplement 1.**
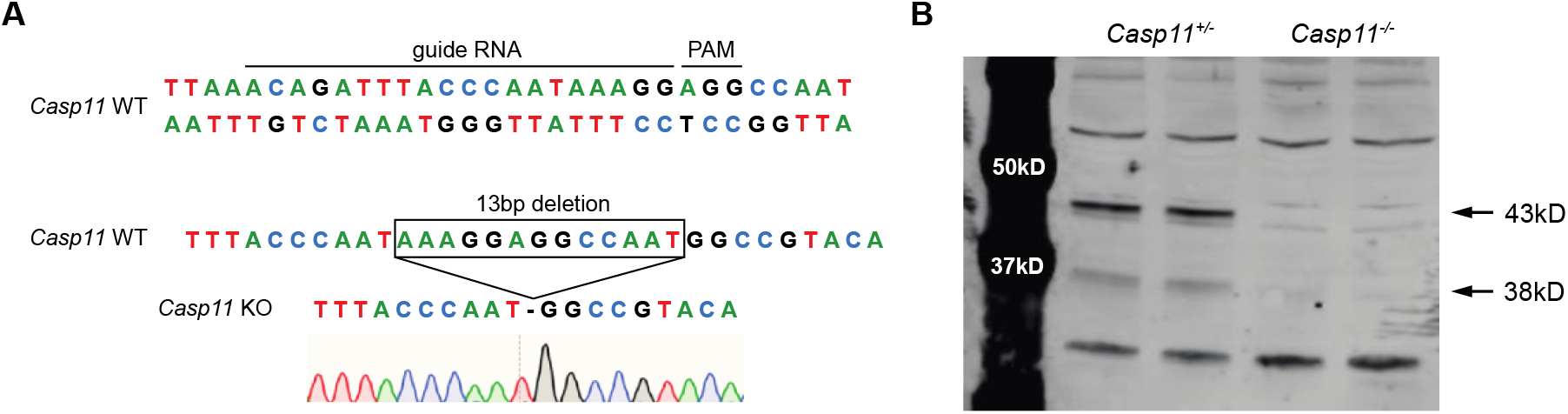
B6.*Nlrc4*^*–/–*^*Casp11*^*–/–*^ mice have a 13bp deletion in *Casp11* and loss of CASP11 protein. (**A**) Endogenous B6.*Casp11* gene with guide RNA sequence and PAM site (above) and resulting edited locus (below). Edited mice have a 13bp deletion in *Casp11* that results in a frameshift mutation. (**B**) Western blot of bone marrow derived macrophage lysates from mice edited at *Casp11* that are either heterozygous (left two lanes) or homozygous knockout (right two lanes). The absence of bands at ∼38 kD and ∼43 kD indicates loss of Caspase-11 protein in the *Casp11*^***–/–***^ mice.

**Figure 2 — figure supplement 2.**
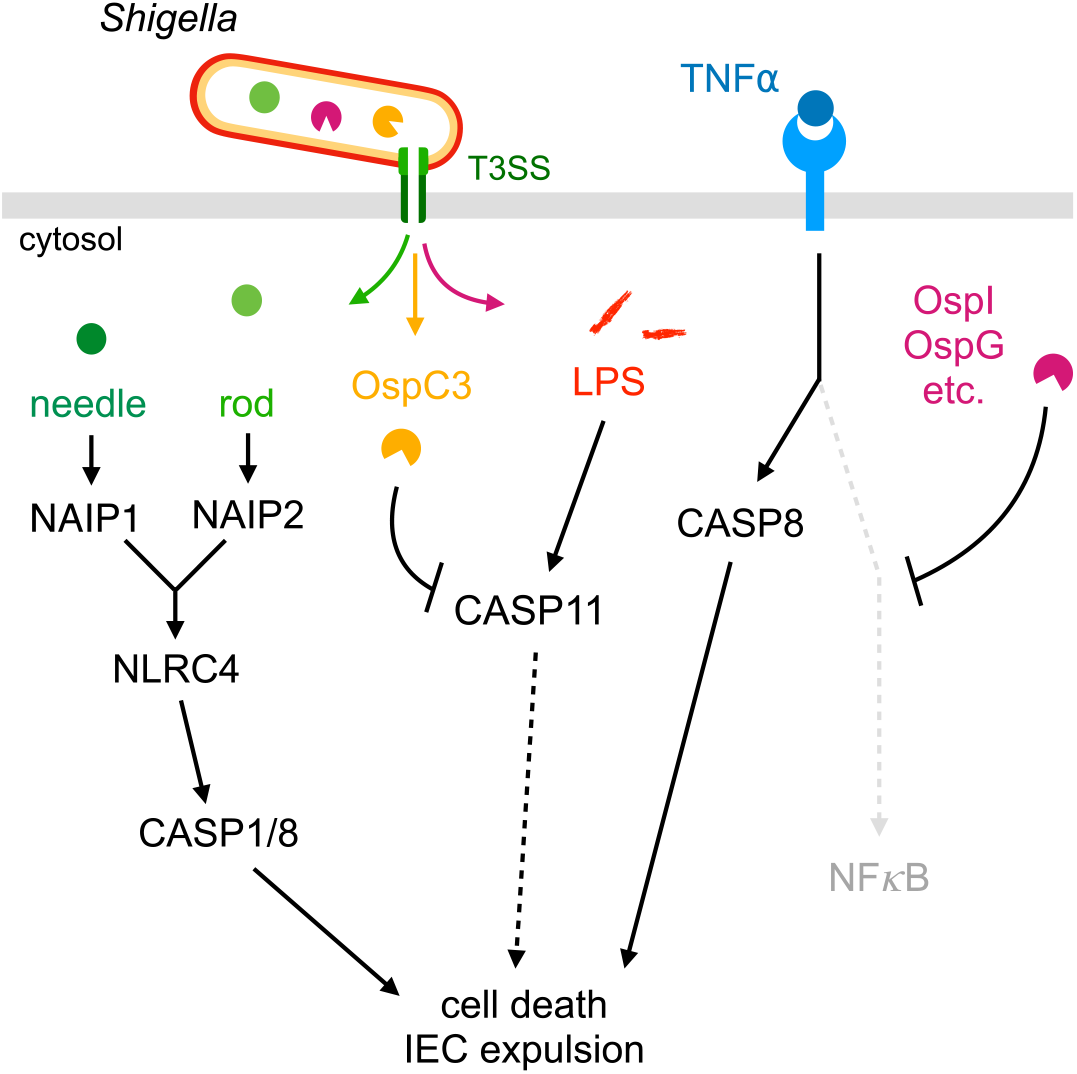
*Shigella* activates mouse cell death pathways. Mouse NAIP–NLRC4, CASP11, and CASP8 respond to *Shigella* pathogen associated molecular patterns or activities to initiate cell death. NAIP1 and NAIP2 receptors bind cytosolic needle and rod proteins (green), respectively, that are secreted through the *Shigella* type three secretion system (T3SS) leading to NLRC4 inflammasome formation and CASP1- or CASP8-dependent pyroptosis. CASP11 recognizes cytosolic *Shigella* LPS (read), leading to non-canonical inflammasome formation and pyroptosis which is partially inhibited when *Shigella* expresses effector OspC3 (yellow). TNFα (blue) initiates CASP8-dependent apoptosis through TNFRI when NF-κB signaling is suppressed by *Shigella* effectors (magenta). All three pathways also lead to cell expulsion when activated in intestinal epithelial cells.

**Figure 7 — figure supplement 1.**
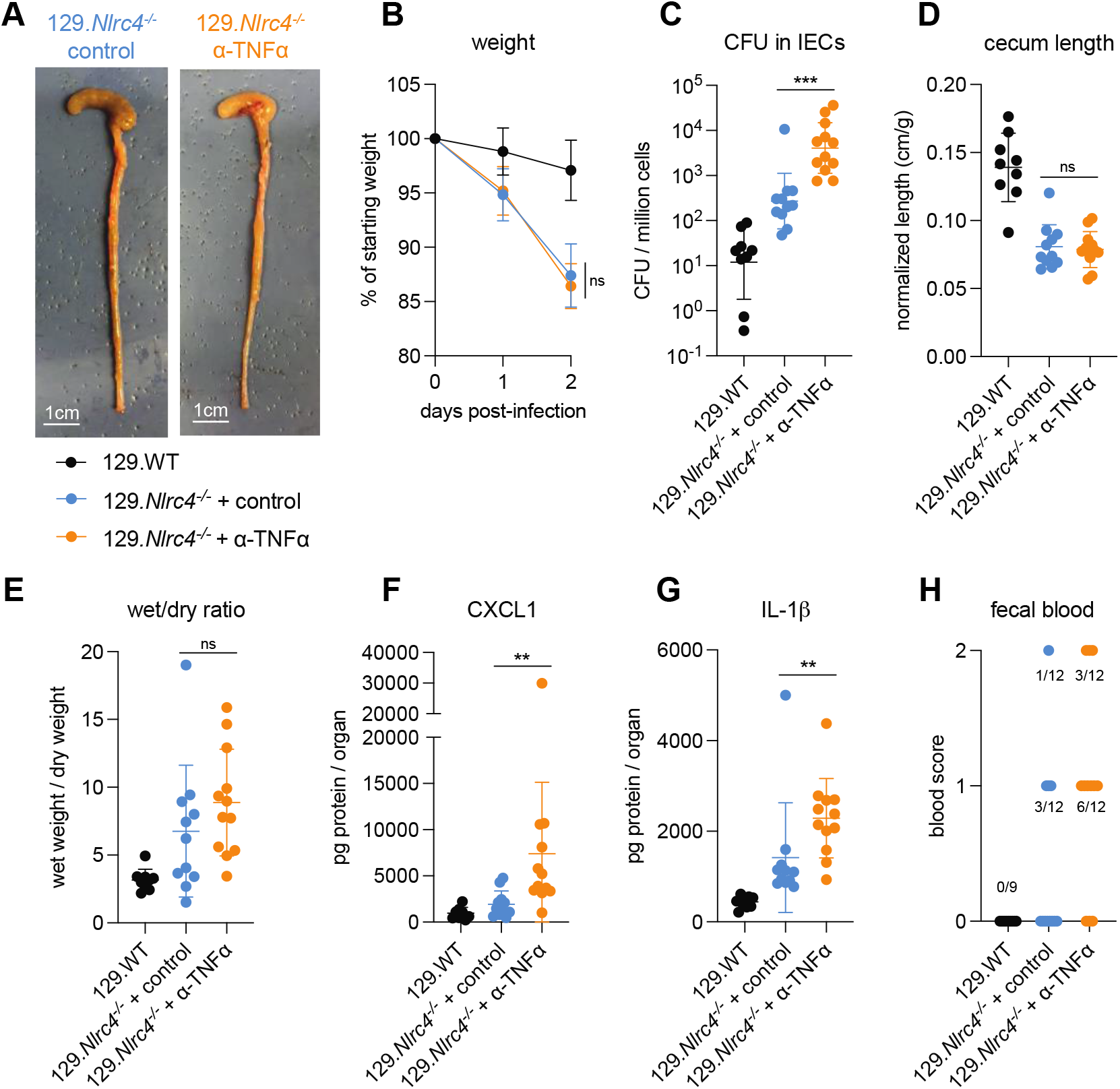
TNFα neutralization renders 129.*Nlrc4*^*–/–*^ mice more susceptible to *Shigella*. (**A-H** 129.WT (black) and 129.*Nlrc4*^***–/–***^ mice were treated orally with 25 mg streptomycin sulfate in water and orally challenged the next day with 10^7^ CFU of WT *Shigella flexneri*. 129.*Nlrc4*^***–/–***^ mice also received 200 μg of either TNFα neutralizing antibody (orange) or isotype control antibody (blue) by intraperitoneal injection daily from one day before infection through sacrifice at two days post-infection. (**A**) Representative images of the cecum and colon from 129.*Nlrc4*^***–/–***^ mice receiving either isotype control or TNFα neutralizing antibody. (**B**) Mouse weights from 0 through 2 days post-infection. Each symbol represents the mean for all mice of the indicated group. (**C**) *Shigella* colony forming units (CFU) per million cells from the combined intestinal epithelial cell (IEC) enriched fraction of gentamicin-treated cecum and colon tissue. (**D**) Quantification of cecum lengths normalized to mouse weight prior to infection; cecum length (cm) / mouse weight (g). (**E**) The ratio of fecal pellet weight when wet (fresh) divided by the fecal pellet weight after overnight drying. Pellets were collected at day two post-infection. (**F, G**) CXCL1 and IL-1β levels measured by ELISA from homogenized cecum and colon tissue of infected mice. (**H**) Blood scores from feces collected at two days post-infection. 1 = occult blood, 2 = macroscopic blood. (**C-H**) Each symbol represents one mouse. Data collected from two independent experiments. Mean ± SD is shown in (**B, D-G**). Geometric mean ± SD is shown in (**C**). Mann-Whitney test, *p < 0.05, **p < 0.01, ***p < 0.001, ****p < 0.0001, ns = not significant (p > 0.05).

